# Chemotherapy-treated breast cancer cells activate the Wnt signaling pathway to enter a diapause-DTP state

**DOI:** 10.1101/2024.03.08.584051

**Authors:** Youssef El Laithy, Willy Antoni Abreu De Oliveira, Anirudh Pabba, Alessandra Qualizza, François Richard, Paraskevi Athanasouli, Carla Rios Luci, Wout De Wispelaere, Larissa Mourao, Siân Hamer, Stijn Moens, Anchel De Jaime-Soguero, Maria Francesca Baietti, Stefan J Hutten, Jos Jonkers, Stephen-John Sammut, Stefaan Soenen, Colinda LGJ Scheele, Alejandra Bruna, Christine Desmedt, Daniela Annibali, Frederic Lluis

## Abstract

The efficacy of chemotherapy is often hindered by the enrichment of a population of cancer cells that enter a drug-tolerant persister (DTP) state, mimicking embryonic diapause, yet the underlying mechanisms of this transition remain poorly understood. This study demonstrates that both parental and chemotherapy-induced Wnt-active (Wnt^High^) cells in Triple-negative breast cancer exhibit transcriptional and functional properties characteristic of DTP cells, including a diapause transcriptional signature, reduced MYC expression, reversible restricted proliferation, and pronounced chemoresistance. Our findings reveal that the *de novo* activation of the Wnt signaling pathway, triggered by the transcriptional upregulation of components essential for canonical Wnt ligand-secretion and -activation, is critical for enriching the diapause-DTP (DTP**^Diap^**) population across various chemotherapy regimens. The diapause-DTP/Wnt^High^ population can be selectively ablated by concomitant, rather than sequential, pharmacological inhibition of Wnt ligand-secretion alongside chemotherapy, highlighting new vulnerabilities in DTP**^Diap^** cell-emergence and potentially yielding a therapeutic opportunity against DTPs. This study shows that activation of Wnt signaling pathway is sufficient and necessary for the induction of a DTP**^Diap^** state and enhances our understanding of the introductory mechanisms driving DTP cell-enrichment upon chemotherapy.

## INTRODUCTION

Accumulating evidence indicates that cancer cells can enter a reversible drug-tolerant persister (DTP) cell-state to evade chemotherapy-induced cell death, leading to incomplete therapy responses and/or recurrence^1–3^. Eradicating DTP cells, or preferably, preventing their formation during cancer treatment, represents a potential strategy to increase cancer sensitization to treatment and, ultimately, improve patient survival rates.

The concept of persisters originates from microbial literature, where it is well established that antibiotic treatment can reduce bacterial burden but sometimes fails to eliminate refractory bacteria^4,5^. Similar subpopulations of DTP cells have been identified in cancer cell lines that survive lethal dosages of targeted therapies or chemotherapy, namely by entering a slow-cycling state. These DTP cells exhibit a distinct transcriptomic profile reminiscent of embryonic diapause, a cellular state used across the animal kingdom, including some mammals, to survive stressful environments^6–8^. In mouse embryonic stem cells (ESC), genetic depletion of c-Myc and n-Myc induces a pluripotent dormant state mimicking diapause^9,10^. Similarly, in cancer cells, pharmacologic inhibition or depletion of MYC promotes a diapause-like state characterized by reduced proliferation and increased resistance to therapy, highlighting the central role of MYC in this process^11^. In agreement, cancer DTP cells negatively correlate with transcriptional MYC hallmark expression. Although the origin of DTP cells through non-genetic processes is well documented, the molecular mechanisms preceding and consequently driving the acquisition of the diapause-DTP (DTP**^Diap^**) cell-state, are still obscure.

DTP**^Diap^** cells exhibit features of epithelial-mesenchymal transition (EMT), which are associated with poor drug responsiveness, a senescence-like gene signature, and enhanced stemness^12–15^. Furthermore, upon discontinuation of treatment—commonly referred to as a drug holiday—DTP cells resume growth and proliferation, and its progeny retains sensitivity to chemotherapy^16^.

Many patients with triple-negative breast cancer (TNBC) initially benefit from preoperative (neoadjuvant) chemotherapy (NAC); however, about 30%–50% develop resistance, leading to poor overall survival rates^17,18^. Drug resistance has conventionally been attributed to the selection of pre-existing resistant (stem) cell populations (intrinsic or Darwinian selection)^19–21^. However, recent research using genomic and transcriptomic deep sequencing of matched longitudinal (pre- and post-NAC treatment) TNBC patient and patient-derived xenograft (PDX) samples has also highlighted the role of acquired (drug-induced) resistance during chemotherapy^22,23^. Interestingly, residual TNBC tumors treated with NAC do not exhibit an enrichment of a breast cancer stem cell (BCSC) population (CD24**^Low^**/CD24**^High^**cells)^23^. DTP cell-enrichment has been demonstrated across distinct chemotherapeutic agents; however, it remains unknown whether the emergence of a DTP**^Diap^** cell-state converges on common downstream molecular mechanisms, even when induced by distinct chemotherapeutic agents with divergent pro-apoptotic mechanisms of action.

Identifying the initial, non-genetic events responsible for the enrichment of DTP**^Diap^** populations in TNBC could pave the way for targeting DTP cells even before their emergence. In this study, we show that *de novo* Wnt transcriptional-activation precedes DTP cell-enrichment, regardless of the chemotherapeutic agent used (docetaxel or carboplatin). This activation of the Wnt signaling pathway by cytotoxic treatment was not limited to *in vitro* 2D-cultured TNBC cell lines but was also consistently observed in 3D-cultured TNBC patient-derived organoid (PDO) models and *in vivo* xenograft models. Our transcriptional and functional analysis of the Wnt-active (Wnt^High^) cell population in parental untreated (UNT) and early-treated samples reveal that Wnt^High^ cells correlate transcriptionally and functionally with DTP**^Diap^** cells. Importantly, activation of the Wnt signaling pathway in parental cells replicates the functional DTP**^Diap^** cell-features, demonstrating that Wnt-activation alone suffices to induce a slow-proliferating DTP cell-state, even under non-chemotherapeutic conditions.

We find that diapause-DTP/Wnt^High^ (DTP**^Diap^/**Wnt**^H^**) cell-enrichment during chemotherapy is driven by increased expression of Wnt ligands, R-spondins, and molecules involved in Wnt ligand-secretion. Surprisingly, although pharmacological inhibition of Wnt ligand-secretion significantly reduces the percentage of the parental DTP**^Diap^/**Wnt**^H^**population, it does not mitigate DTP cell-enrichment once chemotherapy is applied. In contrast, concomitant inhibition of Wnt ligand-secretion and chemotherapy treatment significantly reduces the DTP**^Diap^** population and sensitizes TNBC cell lines, xenograft models, and PDO models to chemotherapy.

Our results demonstrate that distinct therapeutic agents converge on Wnt signaling pathway-activation to drive *de novo* DTP**^Diap^**cell-emergence in various TNBC models. This suggests that a combinatorial treatment strategy involving both Wnt ligand secretion-inhibition and chemotherapy might effectively target the initial mechanisms involved in the enrichment and acquisition of a DTP**^Diap^** cell-phenotype, subsequently benefiting patients with TNBC undergoing systemic chemotherapy.

## RESULTS

### Wnt transcriptional-activation precedes early drug-tolerant cell(s) enrichment upon chemotherapeutic treatment

To study the molecular basis underlying the emergence or enrichment of drug-tolerant cells during therapy, we recapitulated this phenomenon *in vitro*. Three different TNBC cell lines (MDA-MB-231, MDA-MB-468, and PDC-BRC-101) were subjected to two distinct cytotoxic chemotherapeutic agents: docetaxel (DOC) and carboplatin (CAR). These agents operate through different mechanisms of action, with DOC promoting microtubule stabilization and preventing depolymerization, and CAR inducing DNA damage^24,25^. IC50 concentrations (at 72h) were determined for each chemotherapeutic agent for every cell line and used in successive studies (Supplementary Fig. S1A-F). To assess changes in the frequency of putative drug-tolerant cells under chemotherapeutic pressure, the enzymatic activity of aldehyde dehydrogenase 1 (ALDH1), a functional marker in solid tumors for drug resistance and the DTP cell-phenotype in TNBC, was measured^26^. Treatment of all TNBC cell lines with IC50 concentrations of either DOC or CAR led to a significant increase in the levels of ALDH**^High^** cells compared to UNT conditions at 96hours(h) but not at 48h (Fig. 1A), indicating a time-dependent effect on drug-tolerant cell(s) enrichment following chemotherapeutic treatment. Although we detected a small proportion of BCSCs (CD24**^Low^**/CD44**^High^**) population, we recorded no significant enrichment at either 48h or 96h of treatment (Fig. 1B), supporting previous studiese^23^ and suggesting that chemotherapy preferentially enriches for drug-tolerant cells rather than BCSCs during early treatment events.

**Fig. 1:**
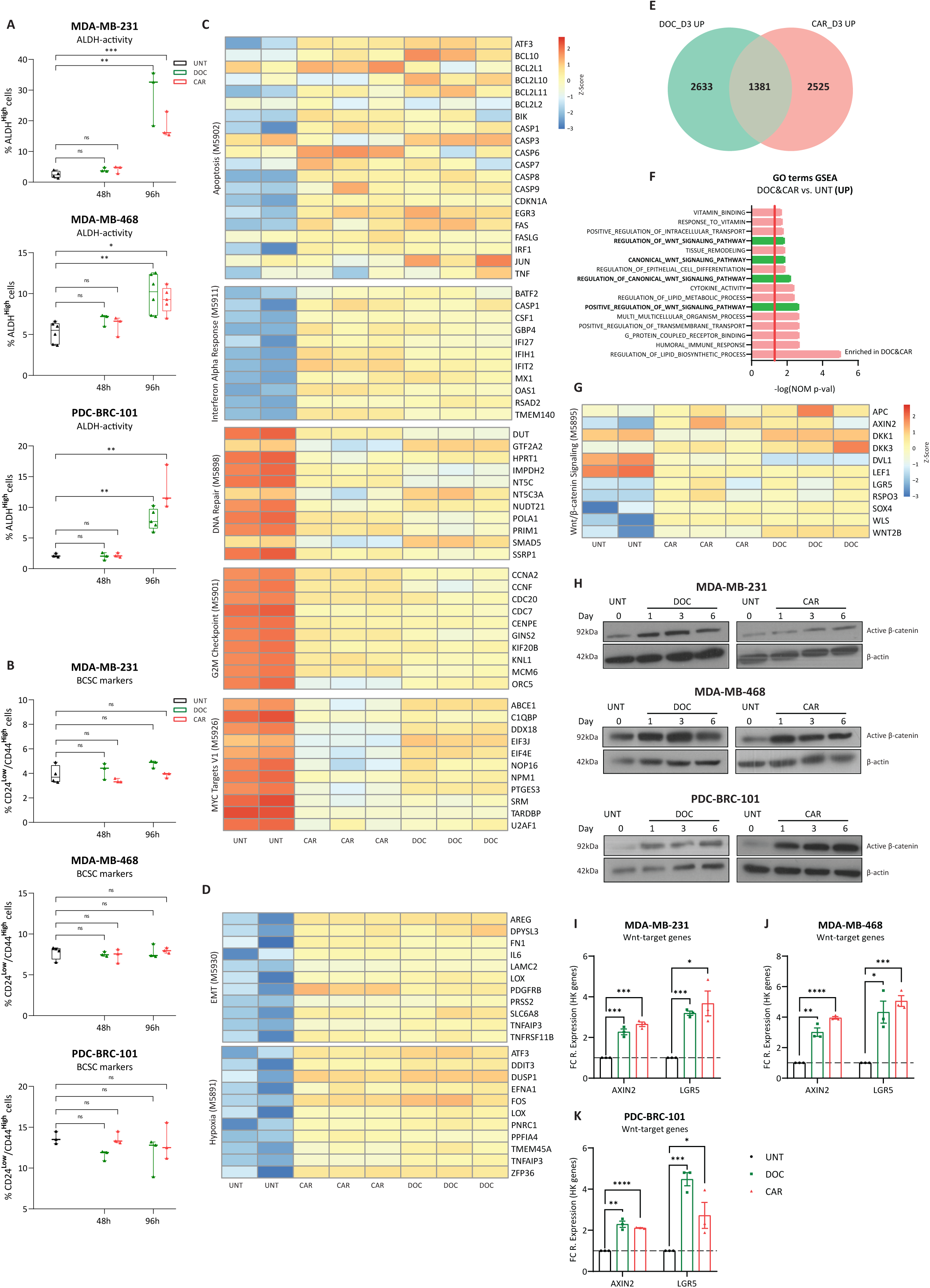
Wnt transcriptional-activation precedes early drug-tolerant cell(s) enrichment upon chemotherapeutic treatment. **A)** Flow cytometry analysis displaying %ALDH**^High^** cells of MDA-MB-231, MDA-MB-468, and PDC-BRC-101 TNBC cell lines treated with DOC (4.5nM) or CAR (35uM, 25uM, 35uM for each cell line, respectively) for 48h and 96h. Multiple t-tests corrected for multiple comparisons using the Holms-Sidak method (n = 3 independent experiments). All data points shown from min. to max. (box and whiskers). **B)** Flow cytometry analysis displaying %CD24^Low^/CD44^High^ cells of MDA-MB-231, MDA-MB-468, and PDC-BRC-101 TNBC cell lines treated with DOC or CAR for 48h and 96h. Multiple t-tests corrected for multiple comparisons using the Holms-Sidak method (n = 3 independent experiments). All data points shown from min. to max. (box and whiskers). **C-D)** Normalized gene expression heatmaps displaying (≈12-20) selected genes (based on DEGs predicting each representative process and/or hallmark) representing different processes and signaling cascades deregulated (relative to UNT cells) in MDA-MB-231 cell line treated with DOC or CAR for 72h. The processes (and subsequent genes selected to represent every process) shown in panels **C-D** were based on enriched gene sets from Hallmarks databases analyzed by one-tailed GSEA ranked by Normalized Enrichment Score (NES) - **Supplementary Fig. S1J, K**. **E)** Venn diagram indicating commonly upregulated (1381) genes between DOC (2633) and CAR (2525) treatment (vs. UNT). **F)** Enriched processes from Gene Ontology (GO) terms databases analyzed by one-tailed GSEA ranked by a positive NES and based on commonly upregulated genes between DOC and CAR treatment. Red line indicates significance threshold value, (-log(NOM p-val) = 1.3) and highlighted bars indicate GO terms linked to Wnt/β-catenin signaling regulation. **G)** Normalized gene counts heatmap displaying (11) selected genes representing Wnt/β-catenin signaling in MDA-MB-231 cell line treated with DOC and CAR for 72h. Genes selected for heatmap **G** were based on Wnt/β-catenin signaling Hallmark – GSEA. Data used to generate panels **C-G** was obtained from mRNA-sequencing of MDA-MB-231 cell line treated with DOC or CAR for 72h). **H)** Western blot analysis of active- (non-phosphorylated) β-catenin in MDA-MB-231, MDA-MB-468, and PDC-BRC-101 cell lines treated with DOC or CAR for 1, 3, or 6 days. **I-K)** Gene expression levels obtained via RT-qPCR of Wnt-target genes (*AXIN2* and *LGR5*) for TNBC cell lines treated with DOC or CAR for 72h, displayed as fold change (to UNT) of 2^-dCt^ values (relative to HK-genes). Multiple t-tests corrected for multiple comparisons using the Holms-Sidak method (n = 3 independent experiments). p values are indicated as *p < 0.05, **p < 0.01, ***p < 0.001, ***p < 0.0001, and ns, not significant.

To gain insights into the signaling cues preceding and, potentially, driving this chemotherapy-mediated enrichment of drug-tolerant (ALDH**^High^**) cells at 96h, we performed bulk transcriptomic analysis of mRNA-sequenced samples obtained from viable/drug-tolerant (DAPI^-^) MDA-MB-231 cells treated with either DOC or CAR at 72h (Supplementary Fig. S1G). Gene Set Enrichment Analysis^27^ (GSEA) using MSigDB^28^ datasets on differentially expressed genes (DEGs – Supplementary Fig. S1H, I, and Supplementary Table 1) between DOC vs. UNT or CAR vs. UNT (FC > 1.5, p-val ≤ 0.05) identified an array of Hallmarks significantly enriched in DOC or CAR treatments, including Apoptosis, p53 Pathway, and Interferon Gamma Response, all of which align with the expected cell stress induced by chemotherapeutic exposure^29,30^ (Fig. 1C, Supplementary Fig. S1J, K, and Supplementary Table 2). Conversely, Hallmarks associated with cell cycle regulation, such as G2M checkpoint, DNA Repair, MYC targets, and E2F targets were significantly downregulated in DOC- and CAR-treatment^31,32^ (Fig. 1C and Supplementary Fig. S1L, M). Interestingly, genetic signatures and processes such as EMT and Hypoxia, both associated with tumorigenesis, chemoresistance, and the DTP cell-phenotype were enriched in response to chemotherapeutic exposure^33,34^ (Fig. 1D and Supplementary Table 2).

To elucidate common enriched transcriptomic alterations among distinct chemotherapeutic agents, we performed Gene Ontology (GO) analysis using the commonly (DOC & CAR vs. UNT) upregulated (1381) genes shared between both drugs (Fig. 1E and Supplementary Tables 1, 2). GO analysis repeatedly highlighted a significant enrichment of (positive) regulation of Canonical Wnt signaling, which was corroborated with an enrichment in the expression of Wnt-targets (*AXIN2* and *LGR5*) and upstream regulators and activators (*WLS* and *WNT2B*) of the pathway (Fig. 1F, G). Conversely, GO analysis using commonly downregulated (785) genes shared between both drugs highlighted significant enrichment in processes related to cell cycle regulation and progression (Supplementary Fig. S1N, O).

Western Blot analysis of active (non-phosphorylated) β-catenin in TNBC cell lines confirmed increased levels of active β-catenin, as early as 24h post-treatment, and prolonged for up to 6 days (Fig. 1H). Consequently, the expression levels of Wnt signaling target-genes (*AXIN2* and *LGR5*) showed a significant increase, confirming the transcriptional activation of the Wnt signaling pathway upon chemotherapeutic treatment (Fig. 1I-K).

Our data reveals an upregulation of canonical Wnt signaling activity preceding DTP cell-enrichment. Notably, this enrichment or activation of the Wnt signaling pathway appears to be a common phenomenon shared among various TNBC cell lines and in response to two distinct cytotoxic treatments.

### Parental and chemotherapy-treated Wnt^High^ cells display DTP^Diap^ cell-properties

To gain a more comprehensive understanding of the transcriptional and functional features of Wnt-active (Wnt^High^) population, we generated clonal MDA-MB-231, MDA-MB-468, and PDC-BRC-101 TNBC cell lines carrying a stable integrated Wnt-transcriptional (β-catenin-TCF/LEF-mediated transcriptional activity) reporter (TOP-GFP/TGP cell lines)^35^ (Fig. 2A). We observed a range of TOP-GFP expression patterns in all three TNBC cell lines cultured in basal (UNT) conditions, with an approximate 6%, 2.5%, and 10% GFP^+^ (referred to hereafter as Wnt^High^) cells in MDA-MB-231-TGP, MDA-MB-468-TGP, and PDC-BRC-101-TGP cell lines, respectively (Fig. 2B – black bars). Upon exposure to either DOC or CAR agents, we observed a significant increase both in the percentage of Wnt^High^ cells in viable/drug-tolerant (DAPI^-^) cells and in the levels/degree of transcriptional Wnt-activation compared to UNT conditions (Fig. 2B and Supplementary Fig. S2A, B). Prolonged exposure to either therapeutic agent for 6 days maintained or increased the percentage of transcriptional Wnt^High^ cells (Fig. 2B), further confirming that the enrichment and/or induction of Wnt^High^ cells is one of the early events directing drug-tolerance.

**Fig. 2:**
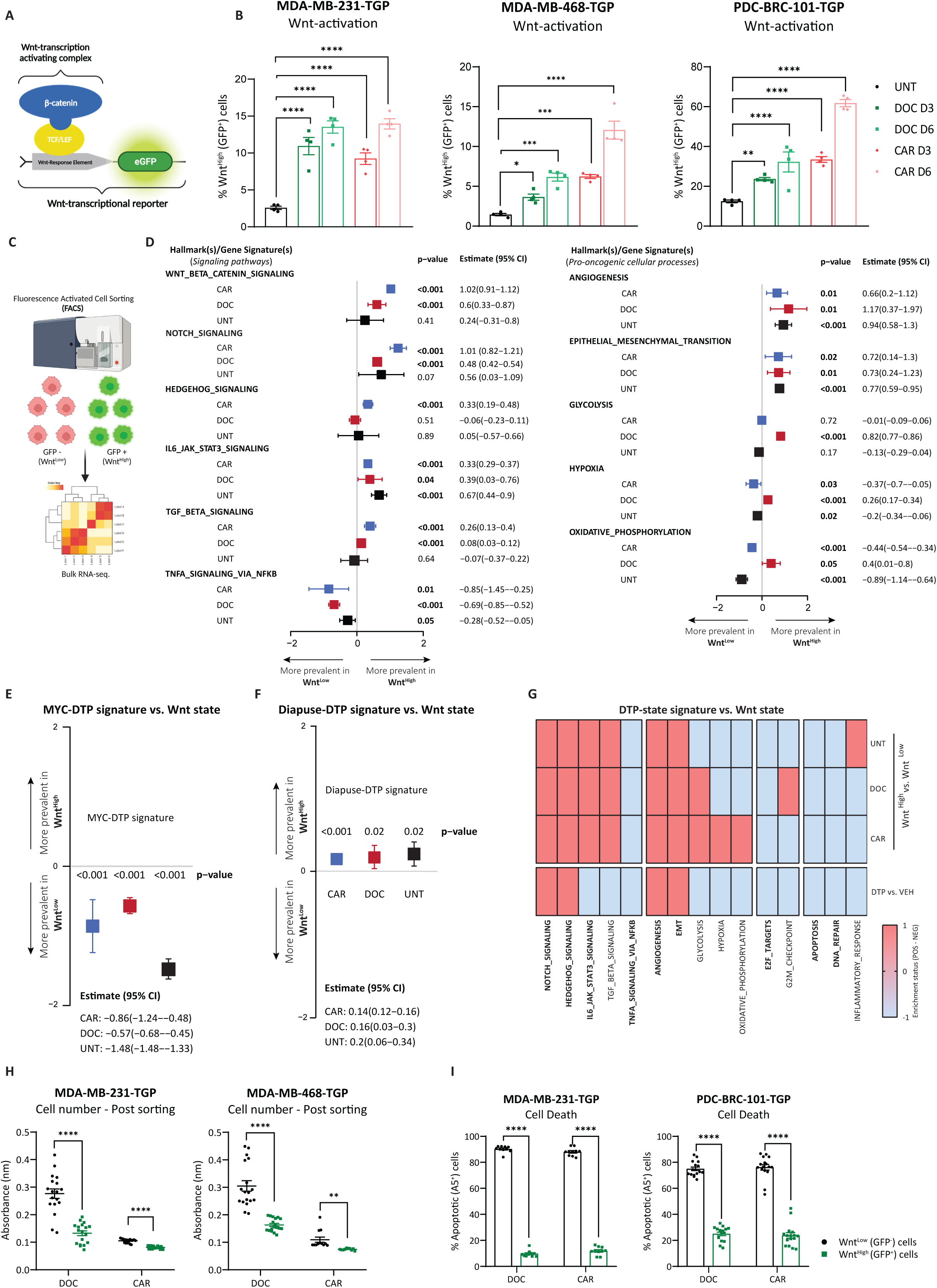
Parental and chemotherapy-treated Wnt^High^ cells display DTP^Diap^ cell-properties. **A)** Schematic representation of Wnt/β-catenin-transcriptional reporter (TOPGFP/TGP). **B)** Flow Cytometry analysis displaying % of Wnt^High^ (GFP^+^) cells for MDA-MB-231-TGP, MDA-MB-468-TGP, and PDC-BRC-101-TGP cell lines treated with DOC or CAR for 3 and 6 days. One-way ANOVA corrected for multiple comparisons using the Holms-Sidak method (n = 4 independent experiments). Data are presented as Mean ± SEM. **C)** Schematic repreentation of the experimental setup designed for studying the transcriptional differences between Wnt^High^ vs. Wnt^Low^ cells. **D)** Forest plots depicting the association between gene signatures and Wnt-status of sorted Wnt^High^ vs. Wnt^Low^ cells obtained from samples treated with CAR or DOC or UNT samples. Gene signatures consist of in-house gene signatures^87^ and Hallmark gene sets from MSigDB analysed using Gene set variation analysis (GSVA). Quartile regression was used to observe the median change in rescaled gene signatures after accounting for batch effects. Signatures having a non-zero positive estimate indicate increased activity in Wnt^High^ cells. **E-F)** Forest plot depicting the association between a MYC Hallmark signature (left) and a DTP**^Diap^** signature^7^ (right) vs. Wnt-status of sorted Wnt^High^ vs. Wnt^Low^ cells obtained from samples treated with CAR or DOC and UNT samples. Analysis performed as in **D**. **G)** Correlation analysis between transcriptional DTP cells^11^ and Wnt-status of sorted Wnt^High^ vs. Wnt^Low^ cells obtained from samples treated with CAR or DOC and UNT samples. Enrichment status score of −1 (light blue) indicates that a given hallmark/process is downregulated in DTPs or sorted Wnt^High^ cells while a score of 1 (red) indicates that a given hallmark/process is upregulated in DTPs or sorted Wnt^High^ cells. **H)** Absorbance values displaying cellular metabolic activity indicating cell number in sorted MDA-MB-231-TGP and MDA-MB-468-TGP cell lines 1 week after sorting (initial treatment before sorting was with DOC or CAR – 72h). Multiple t-tests corrected for multiple comparisons using the Holms-Sidak method (n = 3 independent experiments). Data are presented as Mean ± SEM. **I)** Flow cytometry analysis displaying % of Apoptotic (Annexin V⁺) and their respective Wnt-status (% of Wnt^High^ (GFP⁺) cells) of MDA-MB-231-TGP and PDC-BRC-101-TGP cell lines treated with DOC or CAR for 96h. Multiple t-tests corrected for multiple comparisons using the Holms-Sidak method (n = 5 independent experiments). Data are presented as Mean ± SEM. p values are indicated as *p < 0.05, **p < 0.01, ***p < 0.001, ****p < 0.0001, and ns, not significant.

To unravel the transcriptional discrepancies between the Wnt^Low^ and Wnt^High^ populations, we conducted bulk transcriptomic analysis on mRNA-sequenced samples derived from viable/drug-tolerant (DAPI^-^) MDA-MB-231-dTGP sorted populations under chemotherapy-treated conditions^36^ (Fig. 2C). Gene set variation analysis (GSVA) revealed a plethora of MSigDB Hallmark signatures from DEGs between CAR or DOC (FC > 1.5, p-val ≤ 0.05) in Wnt^High^ vs. Wnt^Low^ sorted cells that were differentially up- or down-regulated (Supplementary Fig. S2C, D, and Supplementary Tables 3-5).

As expected, the Wnt signaling pathway was significantly and positively associated in CAR- and DOC-sorted Wnt^High^ populations compared to Wnt^Low^ populations (Fig. 2D and Supplementary Fig. S2E). While a few signatures exhibited drug-dependent associations, most differentially regulated hallmarks followed a similar associative trend among sorted chemo-treated Wnt^High^ and Wnt^Low^ cells observed across both drug treatments (Fig. 2D and Supplementary Fig. S2E). Notably, developmental signaling pathways, including Hedgehog, Notch, IL-6/JAK/STAT3, and TGF-β signaling, along with hallmarks linked to tumor progression, stemness capacity, and metastasis^37–40^ (e.g., Angiogenesis and EMT), displayed a significant positive association with Wnt^High^ populations in comparison to Wnt^Low^ populations (Fig. 2D and Supplementary Fig. S2E). Conversely, the TNF-α signaling pathway via the NF-κB pathway notably exhibited a significant negative association with Wnt^High^ cells (Fig. 2D and Supplementary Fig. S2E) aligning with previous findings suggesting that active β-catenin can attenuate transcriptional NF-κB activity in breast cancer^41^.

Even though Wnt pathway-activation has been shown to correlate with increased proliferation^42,43^, our transcriptional analysis revealed that Wnt^High^ cells negatively correlated with transcriptional signatures of proliferation, as evidenced by significant reduction of G2M checkpoint and E2F target signatures across chemo-treated Wnt^High^ samples (Supplementary Tables 4, 5).

Recent studies by Rehman et al. and Dhimolea et al. have provided significant insights into the transcriptional landscape of cancer DTP cells, drawing notable parallels with ESCs^7,11^. These works emphasize that DTP cells suppress MYC activity and exhibit a distinctive gene signature associated with embryonic diapause. Interestingly, our transcriptional analyses revealed that chemo-sorted Wnt^High^ cells had a significant negative association with the MYC hallmark signature whilst having a significant positive association with the Rehman et. al. embryonic DTP**^Diap^** gene signature (Fig. 2E, F, and Supplementary Fig. S2E), further providing evidence and highlighting the similarities between the transcriptomes of DTP cells and chemo-sorted Wnt^High^ cells. Furthermore, the coordinated up- and down-regulation of additional hallmarks and processes (Notch, EMT, and Angiogenesis; upregulated) and (E2F targets, DNA repair, and Apoptosis; downregulated) (Fig. 2G) was coequally recorded in DTP cells and chemo-sorted Wnt^High^ cells. We aimed to explore if the observed transcriptional changes in Wnt^High^ cells under chemotherapy pressure were already pre-existing in UNT (i.e. parental cells) conditions (Supplementary Fig. S2F). Despite a positive correlation between Wnt^High^ cells obtained from UNT conditions and the Wnt/β-catenin hallmark signature, the association was not statistically significant (Fig. 2D), consistent with our previous data highlighting lower levels of Wnt signaling intensity in UNT samples (Supplementary Fig. S2B). Interestingly, parental-sorted Wnt^High^ cells exhibited overall positive associations with developmental signaling pathways and significant positive associations with EMT, while exhibiting significant negative associations with cell cycle hallmarks, following a similar trend as chemo-sorted Wnt^High^ cells (Fig. 2D, Supplementary Fig. S2G, and Supplementary Tables 3-5). Moreover, and with respect to DTP cell-phenotype, parental Wnt^High^ cells displayed similar tendencies to chemo-sorted Wnt^High^ cells; presenting a significant negative association with the MYC hallmark signature (Fig. 2E – UNT), a significant positive association with the Rehman et. al. DTP**^Diap^** gene signature (Fig. 2F – UNT), and other matching hallmarks and processes (Fig. 2G – UNT), highlighting transcriptional Wnt-activity as a functional marker for DTP**^Diap^** cells even in chemo-naïve conditions. The significance of these correlations becomes more apparent in chemo-treated conditions, where transcriptional Wnt-activation is strongly exacerbated.

Functional analyses confirmed reduced cell proliferation measured by cell number observed in chemo-sorted Wnt^High^ compared to Wnt^Low^ populations (Fig. 2H). Furthermore, co-staining of GFP-expression (reporter for Wnt-activity) alongside the apoptotic marker Annexin V revealed that Wnt^High^ cells displayed reduced apoptotic activity confirming their enhanced drug-tolerant state (Fig. 2I).

In summary, our comprehensive data demonstrates that Wnt^High^ TNBC cells not only exhibit a transcriptional DTP cell-phenotype but also function effectively as *bona fide* DTP**^Diap^** cells in parental drug-naïve cells or under chemotherapeutic conditions. This suggests that transcriptional Wnt-activity may serve as a distinct functional biomarker of the DTP**^Diap^** cell-state in parental cells, but especially in early response to chemotherapeutic challenges.

### Wnt pathway-activation is sufficient to induce a DTP^Diap^ state in parental TNBC cells

Next, we investigated whether activating the Wnt signaling pathway in parental (chemo-naïve) TNBC cells is sufficient to induce a paused, drug-tolerant state similar to that observed in treatment-induced DTP**^Diap^/**Wnt**^H^**cells.

We stimulated Wnt transcriptional-activation using two distinct GSK3 inhibitors, CHIR99021 (CHIR) and 6-Bromoindirubin-3’-oxime (BIO), which stabilize β-catenin^44^. Treatment of parental MDA-MB-231-TGP and MDA-MB-468-TGP cell lines with either CHIR or BIO resulted in activation of the Wnt signaling pathway as evidenced by a significant increase (>90%) in levels of DTP**^Diap^/**Wnt**^H^** cells recorded via FACS (Fig. 3A, B) alongside a significant increase in transcriptional levels of Wnt target-gene *AXIN2* (Supplementary Fig. S3A).

**Fig. 3:**
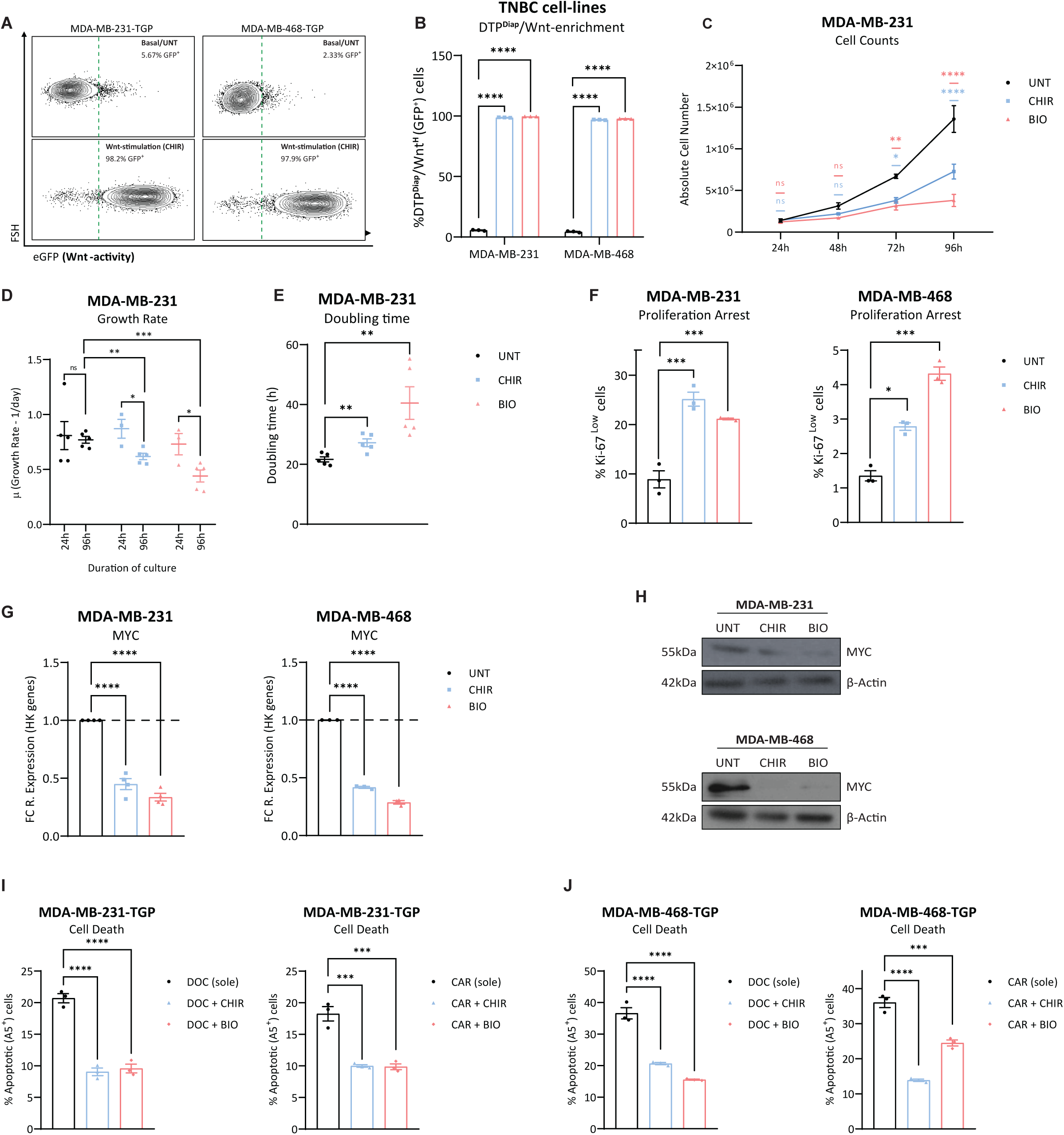
Wnt pathway-activation is sufficient to induce a DTP^Diap^ state in parental TNBC cells. **A)** Representative flow cytometry contour plots displaying % of DTP**^Diap^**/Wnt**^H^** (GFP^+^) cells for MDA-MB-231-TGP and MDA-MB-468-TGP cell lines in basal (UNT) culture conditions (top) and under Wnt-stimulation conditions (CHIR, 8uM, 72h – bottom). Right of the dashed green line indicates DTP**^Diap^**/Wnt**^H^** (GFP^+^) cells. **B)** Flow cytometry analysis displaying % of DTP**^Diap^**/Wnt**^H^**(GFP⁺) cells of MDA-MB-231-TGP and MDA-MB-468-TGP cell lines under Wnt-stimulatory conditions, treated with CHIR (8uM) or BIO (3uM) for 72h. Two-way ANOVA corrected for multiple comparisons using Tukey’s test (n = 3 independent experiments). Data are presented as Mean ± SEM. **C)** Absolute cell number of MDA-MB-231 cell line treated with CHIR or BIO for a time-course of 96h. Two-way ANOVA corrected for multiple comparisons using Tukey’s test (n = 3 independent experiments). Data are presented as Mean ± SEM. **D)** Growth rate of MDA-MB-231 cell line at 24h and 96h under UNT or under CHIR and BIO treatment conditions. Multiple t-tests corrected for multiple comparisons using the Holms-Sidak method (n = 3 independent experiments). Data are represented as Mean ± SEM. **E)** Doubling time of MDA-MB-231 cell line at 96h under UNT or under CHIR and BIO treatment conditions. Multiple t-tests corrected for multiple comparisons using the Holms-Sidak method (n = 3 independent experiments). Data are represented as Mean ± SEM. **F)** Flow cytometry analysis displaying % of cells in proliferation arrest (Ki-67^low^ cells) from MDA-MB-231 and MDA-MB-468 cell lines treated with CHIR or BIO for 96h. Multiple t-tests corrected for multiple comparisons using the Holms-Sidak method (n = 3 independent experiments). Data are presented as Mean ± SEM. **G)** Gene expression levels obtained via RT-qPCR of MYC for MDA-MB-231 and MDA-MB-468 cell lines treated with CHIR or BIO for 72h, displayed as fold change (to UNT) of 2^-dCt^ values (relative to HK-genes). Unpaired t-tests based on relative expression values (2^-dCt^) (n = 3 independent experiments). Data are presented as Mean ± SEM. **H)** Western blot analysis of MYC in MDA-MB-231 and MDA-MB-468 cell lines treated with CHIR or BIO for 72h. **I-J)** Flow cytometry analysis displaying % of Annexin V⁺ cells of MDA-MB-231-TGP (left) and MDA-MB-468-TGP (right) cell lines treated with DOC or CAR for 72h (sole or pre-treated with CHIR or BIO for 48h). One-way ANOVA corrected for multiple comparisons using the Dunnett method (n = 3 independent experiments). Data are presented as Mean ± SEM. p values are indicated as *p < 0.05, **p < 0.01, ***p < 0.001, ****p < 0.0001, and ns, not significant.

CHIR and BIO treatments were found to significantly inhibit the proliferation of MDA-MB-231 cells, as observed with cell counts recorded over a 96h time-course, compared to UNT cells (Fig. 3C). While UNT MDA-MB-231 cells maintained a consistent growth rate (µ) over the 96h time-course (µ: 0.8 at 24h – 0.77 at 96h), CHIR and BIO treated cells exhibited a significant decrease in growth rate at 96h (µ: 0.61 and 0.44 at 96h for CHIR and BIO, respectively) (Fig. 3D). Moreover, at 96h, the doubling time (27.2h and 40.4h) for CHIR and BIO, respectively, was significantly increased when compared to UNT cells (doubling time = 21.66h) (Fig. 3E). These findings were corroborated by additional testing using the MDA-MB-468 cell line, whereby cell counts (Supplementary Fig. S3B) and growth rate (Supplementary Fig. S3C) were significantly decreased in CHIR and BIO treatment conditions while doubling time (Supplementary Fig. S3D) was significantly increased, mirroring the results obtained using the MDA-MB-231 cell line. Notably, the decline in cell count was not attributed to apoptotic effects exerted via either GSK3 inhibitors, as confirmed by an Annexin V staining (Supplementary Fig. S3E). Further analysis using immunofluorescence-based staining of proliferation marker Ki-67 highlighted a significant induction of growth arrest under Wnt-stimulatory conditions (Fig. 3F), suggesting that activation of the Wnt signaling pathway prompts a state of paused proliferation in TNBC cell lines.

Parallelly, and in conjunction with what has been reported in DTP**^Diap^**cells and in chemo-treated DTP**^Diap^/**Wnt**^H^** cells, treatment with CHIR or BIO resulted in significant decrease of transcriptional *MYC* and *NMYC* expression levels and MYC protein levels in MDA-MB-231 and MDA-MB-468 cell lines (Fig. 3G, H and Supplementary Fig. S3F), highlighting a direct correlation between Wnt-activation and an anti-proliferative and a suppressed MYC phenotype. To evaluate if Wnt-activation could induce a functional drug-tolerant state, we pre-treated TNBC cell lines with CHIR and BIO before administering chemotherapy. The results showed a significant reduction in apoptosis induction in pre-treatment Wnt-stimulated cells compared to those with sole chemotherapy treatment, across both DOC and CAR agents and in both cell lines. (Fig. 3I, J, and Supplementary Fig. S3G).

Contrary to previous findings that associate Wnt-activation with increased proliferation and CSCs^45,46^, our results suggest that, in TNBC, Wnt-activation can promote a paused proliferative state, reduced/suppressed MYC levels, and enhanced chemoresistance – effectively mimicking the chemo-mediated DTP**^Diap^**cell phenotype.

### Induction of transient *de novo* Wnt signaling transcriptional-activation in response to chemotherapy in TNBC cell lines

The evolution of DTP cells during treatment continues to be a hotly debated topic with some studies suggest that a pre-existing DTP-population is simply enriched during therapy^19–21^, while others propose that cells undergo a temporary phenotypic transition due to cellular plasticity^1,22,23^. To better understand the dynamics of this process, we monitored the activation of the Wnt-reporter (TOP-GFP/TGP) in the Wnt^High^ population through live-cell imaging.

While live-cell imaging analysis of the MDA-MB-231-TGP cell line under UNT conditions did not reveal significant changes in the total levels of Wnt^High^ cells (Fig. 4A – black line), treatment with either DOC or CAR resulted in a gradual enrichment of Wnt^High^ cells over a similar chemo-culture timespan (Fig. 4A – green and red lines), consistent with our previous FACS-based results.

**Fig. 4:**
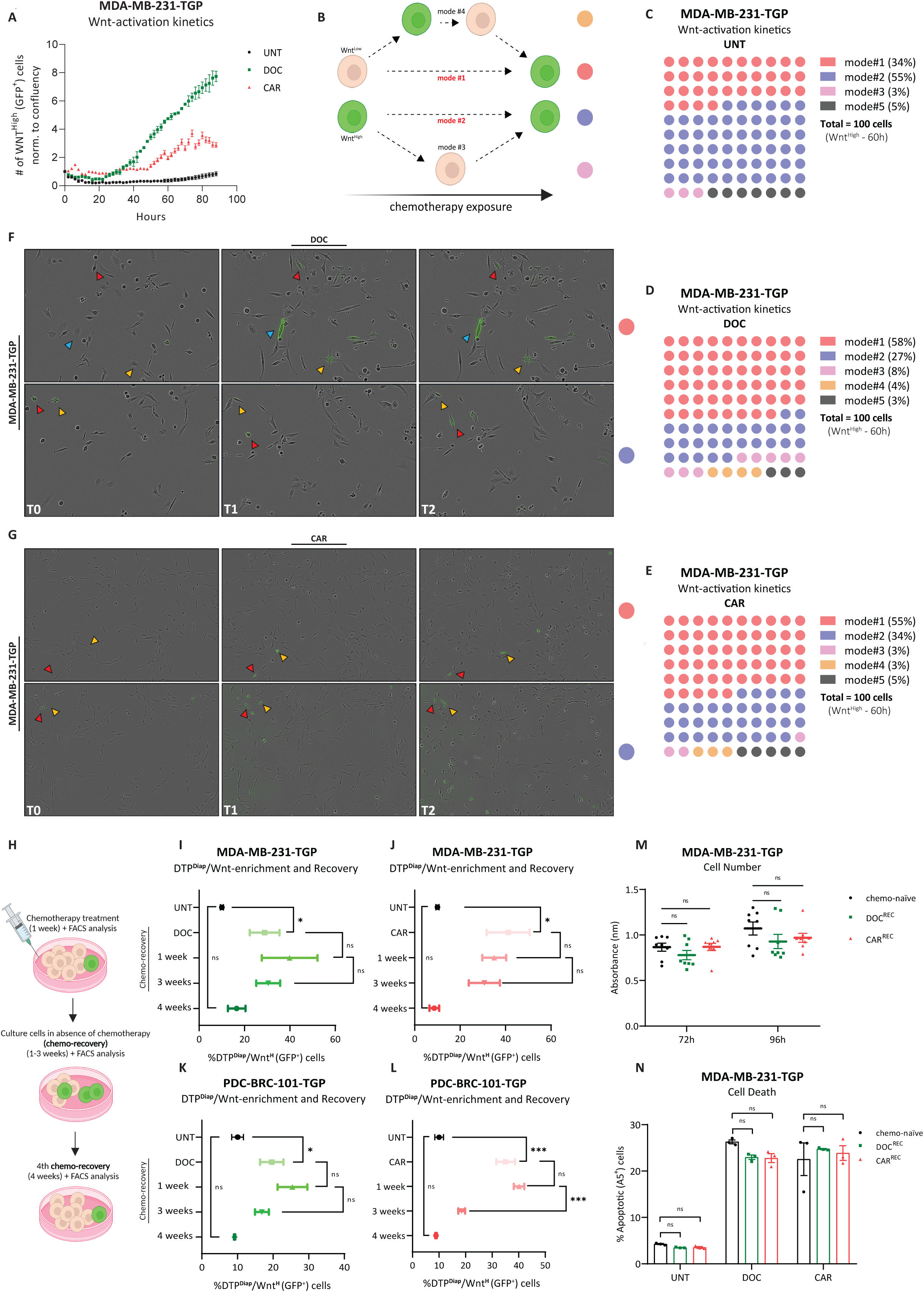
Induction of transient *de novo* Wnt signaling transcriptional-activation in response to chemotherapy in TNBC cell lines. **A)** Number of Wnt^High^ (GFP^+^) cells detected by live-cell imaging normalized to the confluency of the well (total number of cells recorded) for MDA-MB-231-TGP cell line treated with DOC or CAR. **B)** Schematic representation of different fluctuation dynamics of Wnt-activation – mode numbered and color coded. **C-E)** Live-imaging quantification of different possible mechanisms of chemotherapy-induced Wnt-activation in MDA-MB-231-TGP TNBC cell line UNT or treated with DOC or CAR. n = 100 cells tracked every 2h for 60h, per treatment condition. Every single cell tracked is represented as a circle and color coded with the scheme shown in **B**. **F-G)** Snapshots of still-frames from time-lapse live imaging experiments of MDA-MB-231-TGP TNBC cell line treated with DOC (top) or CAR (bottom) presenting mode #1 and mode #2 (color coding in scheme shown in **B**. T0 indicates 0h, T 1 indicates 30h, and T2 indicates approx. 50h. Yellow, red, and blue arrows indicate the same cell followed over the treatment period spanning different images (horizontal). **H)** Schematic representation of the experimental setup for chemotherapy treatment and recovery. **I-L)** Flow Cytometry analysis displaying % of DTP**^Diap^**/Wnt**^H^** (GFP⁺) cells for MDA-MB-231-TGP and PDC-BRC-101-TGP cell lines treated with DOC or CAR for 1 week followed by removal of chemotherapy for 1-, 3-, and 4-weeks post-recovery. Two-tailed Unpaired t-test (n=3 independent experiments). Data are presented as Mean ± SEM. **M)** Metabolic activity levels reflecting cell number and proliferation rates of MDA-MB-231-TGP cell lines (chemo-naïve vs. DOC^REC^ vs. CAR^REC^), all in UNT conditions at 72h and 96h. Two-way ANOVA corrected for multiple comparisons using Tukey’s test (n=3 independent experiments). Data are presented as Mean ± SEM. **N)** Flow cytometry analysis displaying % of Annexin V^+^ cells of MDA-MB-231-TGP cell lines (chemo-naïve vs. DOC^REC^ vs. CAR^REC^), UNT or treated with DOC or CAR for 72h. Two-way ANOVA corrected for multiple comparisons using Tukey’s test (n=3 independent experiments). Data are presented as Mean ± SEM. p values are indicated as *p < 0.05, **p < 0.01, ***p < 0.001, ****p < 0.0001, and ns, not significant.

To visualize Wnt transcriptional-activation dynamics at single-cell resolution, we tracked the original Wnt-state of Wnt^High^ cells (starting at 60h back to 0h) under UNT or chemotherapy-treated conditions, defining different dynamics of Wnt-activation (Fig. 4B). In UNT conditions, 55% of Wnt^High^ cells in the MDA-MB-231-TGP cell line observed at 60h were initially Wnt^High^ at T_0_, while 34% were activated during the culture span (mode #2 and #1, respectively) (Fig. 4C, Supplementary rep. images Fig. S4A, and Supplementary Videos SV1, V2). In contrast, under DOC or CAR treatment conditions, the majority of Wnt^High^ cells at 60h (58% and 55%, respectively) were *de novo*-activated during treatment, indicating chemotherapy-induced Wnt-activation in initially Wnt^Low^ cells (GFP^-^ at T_0_ – mode #1). Conversely, only 27% and 34% of Wnt^High^ cells in DOC and CAR treatment conditions were initially Wnt^High^ at T_0_ (mode #2) (Fig. 4D, E, rep. images Fig. 4F, G, Supplementary rep. images Fig. S4B, and Supplementary Videos SV3-6). These findings suggest that chemotherapy-induced Wnt pathway-enrichment mainly results from *de novo* activation rather than only passive selection of initially Wnt^High^ cells. Additional modes of Wnt-transcriptional activation dynamics were observed in a minority of cases (mode #3 and #4) while cells that fell out of the imaging frame were considered of unknown origin/state (mode #5). In accordance with these findings, similar observations were validated using the PDC-BRC-101-TGP cell line (Supplementary Fig. S4D-G).

Next, we assessed the population dynamics of chemotherapy-induced DTP**^Diap^/**Wnt**^H^** cells after treatment was halted (Fig. 4H). Interestingly, the percentage of DTP**^Diap^/**Wnt**^H^** cells was stabilized or even increased upon chemotherapy removal for up to 3 weeks (Fig. 4I-L). Prolonged culture of these cells for 4 weeks in chemo-free (i.e., a drug-holiday) conditions resulted in a significant reduction in the percentage of DTP**^Diap^/**Wnt**^H^** cells, re-establishing the Wnt-population to levels similar to that of UNT/basal-cultured cells (Fig. 4I-L). This data indicates that the chemotherapy-induced DTP**^Diap^/**Wnt**^H^**phenotype is transient and reversible at the population level once treatment pressure is removed (Fig. 4I-L and Supplementary Fig. S4H). Furthermore, upon restoration of levels of Wnt-activation to that of UNT/basal-cultured cells, we observed that MDA-MB-231-TGP cell lines recovering DOC and CAR treatments (DOC**^REC^**and CAR**^REC^**, respectively) also reverted to normal (i.e., relative to UNT) rates of proliferation and drug tolerance (Fig. 4M, N), highlighting the transient nature of the DTP-phenotype mediated via chemotherapy-induced Wnt-activation.

Altogether, these findings show that the DTP**^Diap^/**Wnt**^H^**phenotype results namely from a *de novo* chemotherapy-driven action rather than solely representing a manifestation of an inherently chemotherapy-resistant subpopulation selected under treatment pressure. Notably, upon chemotherapy removal, Wnt-activity levels revert to baseline levels, indicating a transient enrichment of a DTP**^Diap^/**Wnt**^H^**cell-state dependent on chemotherapy pressure.

### Chemotherapeutic treatment induces elevated transcriptional expression of Wnt ligands, Wnt enhancers, and Wnt secretion machinery components

The Wnt signaling pathway is highly conserved and activated via the binding of (19) extracellular Wnt ligands (Wnts) to membrane receptors^47,48^. Secretion of Wnt ligands requires the action of the acyltransferase Porcupine (PORCN) followed by Wntless/evenness interrupted (WLS/Evi) which supports transport of Wnts from the Trans-Golgi Network to the plasma membrane. In addition, the Rspo protein family has been shown to enhance Wnt ligand activity to further promote Wnt pathway-activation^47,48^. A RT-qPCR screening, focusing on established canonical Wnt ligands (Wnt-1, Wnt-2, Wnt-2b, Wnt-3, Wnt-3a, and Wnt-7b), Wnt ligand enhancers/amplifiers (Rspo1-4), and Wnt secretion machinery components (WLS/Evi and PORCN), revealed that the transcriptional expression of key genes, including *WNT2B*, *WNT3*, *WNT3A*, *WNT7B*, *RSPO1*, *RSPO3*, *WLS*, and *PORCN* was found to be steadily expressed in basal conditions across all analyzed TNBC cell lines (Supplementary Fig. S5A-C).

Importantly, these genes exhibited a consistent and statistically significant increase in expression levels under chemotherapy-treatment conditions (Fig. 5A-C) suggesting that chemotherapeutic exposure actively promotes elevated transcription levels of several key components involved in canonical Wnt pathway-activation.

**Fig. 5:**
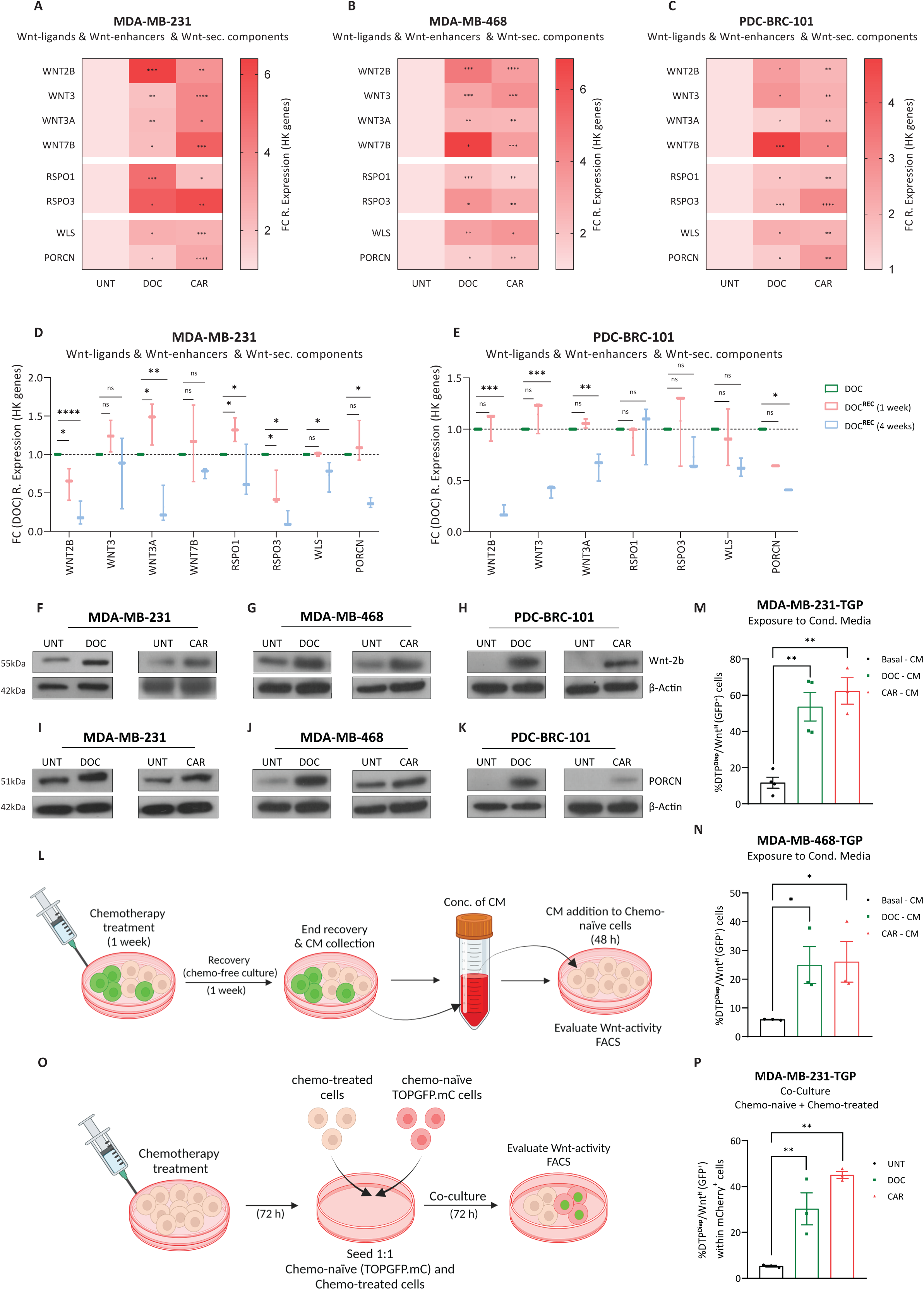
Chemotherapeutic treatment induces elevated transcriptional expression of Wnt ligands, Wnt enhancers, and Wnt secretion machinery components. **A-C)** Heatmaps showing gene expression levels obtained via RT-qPCR of Wnt ligands (WNT2B, WNT3, WNT3A and WNT7B), Wnt enhancers (RSPO1 and RSPO3), and Wnt ligand secretion machinery components (WLS and PORCN) for MDA-MB-231, MDA-MB-468, and PDC-BRC-101 cell lines treated with DOC or CAR for 72h, displayed as fold change (to UNT) of 2^-dCt^ values (relative to HK-genes). Unpaired t-tests based on relative expression values (2^-dCt^) (n = 3 independent experiments). **D-E)** Gene expression levels as in **A-C** for MDA-MB-231 and PDC-BRC-101 cell lines treated with DOC following the treatment scheme shown in Fig. 4H., displayed as fold change (to DOC) of 2^-dCt^ values (relative to HK-genes). Unpaired t-tests based on relative expression values (2dCt) (n = 3 independent experiments). Data are presented as Mean ± SEM. **F-H)** Western blot analysis of Wnt ligand Wnt-2b in MDA-MB-231, MDA-MB-468, and PDC-BRC-101 cell lines treated with DOC or CAR for 72h. **I-K)** Western blot analysis of PORCN in TNBC cell lines treated with DOC or CAR for 72h. **l)** Schematic representation of Conditioned Media (CM) experimental setup. **M-N)** Flow Cytometry analysis displaying % of DTP**^Diap^**/Wnt**^H^**(GFP⁺) cells of MDA-MB-231-TGP and MDA-MB-468-TGP cell lines cultured with concentrated Basal-, DOC-, or CAR-CM for 48h. Unpaired t-tests (n = 3 independent experiments). Data are presented as Mean ± SEM. **O)** Schematic representation of Co-culture experiment setup. **P)** Flow Cytometry analysis displaying % of DTP**^Diap^**/Wnt**^H^** (GFP⁺) cells of chemo-naïve MDA-MB-231-TGP.mC co-cultured with chemo-treated DOC or CAR MDA-MB-231 cells for 72h. Unpaired t-tests (n = 3 independent experiments). Data are presented as Mean ± SEM. p values are indicated as *p < 0.05, **p < 0.01, ***p < 0.001, ****p < 0.0001, and ns, not significant.

Subsequently, following chemotherapy removal and under one week chemo-recovery conditions, the majority of Wnt-activation components maintained elevated expression levels (Fig. 5D, E – pink bars) However, after four weeks of culture in chemo-free conditions and coinciding with the previous results displaying the return of DTP**^Diap^/**Wnt**^H^** cells to basal levels (Fig. 4I-L), we observed a corresponding decrease in expression level patterns of Wnt-activation components (Fig. 5D, E – light blue bars). This correlation between the expression levels of Wnt-activation components and the dynamic induction of a DTP**^Diap^/**Wnt**^H^**population highlights the transient nature of chemotherapy-induced Wnt-activation, providing a possible mechanism for the enrichment of the DTP**^Diap^/**Wnt**^H^** population in response to treatment.

Western blot analysis confirmed the upregulation of the Wnt ligand Wnt-2b and the acyltransferase PORCN, in response to either DOC or CAR treatments across all analyzed TNBC cell lines (Fig. 5F-K).

Treatment of chemo-naïve cells with concentrated conditioned media (CM) derived from MDA-MB-231 and MDA-MB-468 TNBC cell lines (Fig. 5L) confirmed increased and functional presence of Wnt ligands in media collected under one week chemo-recovery conditions, resulting in a significant increase in DTP**^Diap^/**Wnt**^H^** cells when compared to chemo-naïve cells treated with UNT CM (Fig. 5M, N). In a parallel experiment, we co-cultured chemo-naïve (MDA-MB-231-mCherry-TGP) cells with chemo-recovering (MDA-MB-231) cells (Fig. 4O). This co-culturing approach similarly resulted in a substantial and statistically significant increase in the percentage of DTP**^Diap^/**Wnt**^H^**cells within the chemo-naïve cell population (Fig. 5P).

In summary, our findings demonstrate that chemotherapeutic treatment leads to elevated expression levels of Wnt ligands, -enhancers, and -components of the Wnt secretory apparatus. Significantly, concentrated CM obtained from chemo-treated and recovered cells induces an enrichment in the DTP**^Diap^/**Wnt**^H^** cell-population, substantiating the functional impact of the chemo-secreted factors.

### Wnt ligand secretion-inhibition hinders DTP^Diap^/Wnt^H^ population enrichment

The induction and/or enrichment of treatment-persistent residual tumor cells represents an important barrier to curative outcomes. Therefore, a better understanding of the therapeutic vulnerabilities of the DTP**^Diap^/**Wnt**^H^** cell-state potentially has major clinical implications.

We stably transduced MDA-MB-231, MBD-MB-468, and PDC-BRC-101 TNBC cell lines with lentiviral shRNA constructs designed to silence the acetyltransferase PORCN, abrogating the O-palmitoylation and functional secretion of Wnt ligands^49,50^ and resulting in a substantial reduction (approx. 80-90%) in PORCN mRNA levels (Supplementary Fig. S6A-C). PORCN silenced (shPORCN#1) cell lines exposed to either chemotherapeutic agent revealed reduced levels of active β-catenin or percentage of DTP**^Diap^/**Wnt**^H^** cells compared to control (shPLKO) lines, confirming an essential role for PORCN in chemotherapy-induced DTP**^Diap^/**Wnt**^H^**cell-enrichment (Fig. 6A-C and Supplementary Fig. S6D). While the silencing of PORCN had no impact on cell viability in basal (UNT) culture conditions, we observed a marked and significant increase in levels of apoptotic and necrotic (Annexin V^+^ and DAPI^+^) cells in shPORCN#1 TNBC cell lines compared to shPLKO lines upon chemotherapy exposure, indicating a strong sensitization role of PORCN-inhibition (Fig. 6D-F). We validated the reduction of the chemotherapy-induced DTP**^Diap^/**Wnt**^H^**population and increased sensitization to chemotherapy using a second independent lentiviral shRNA construct targeting PORCN (shPORCN#4 – Supplementary Fig. S6E-G) confirming that genetic inhibition of Wnt ligand-secretion effectively hinders the advent of a chemotherapy-induced DTP**^Diap^/**Wnt**^H^**population while significantly enhancing the sensitization of TNBC cell lines to chemotherapy.

**Fig. 6:**
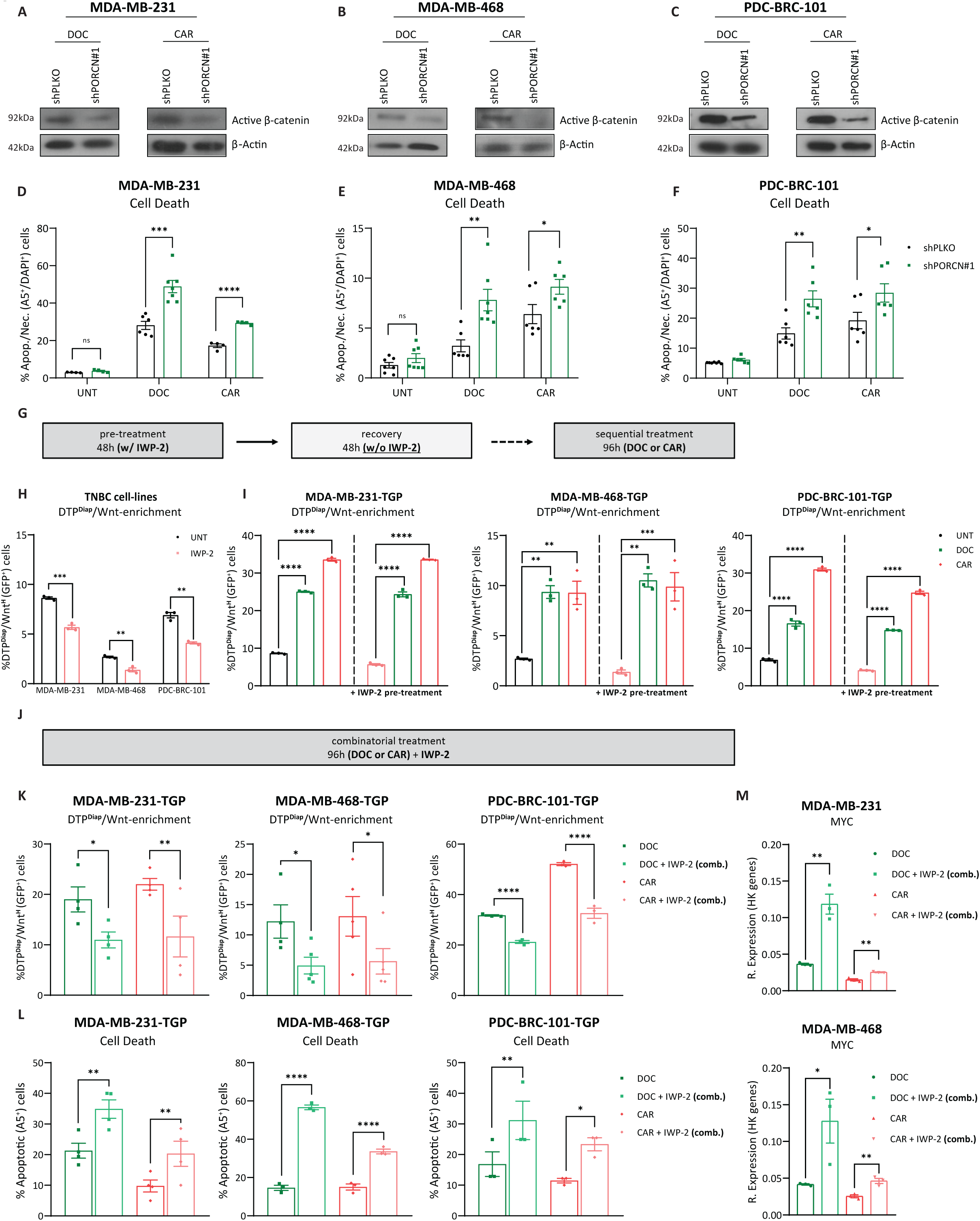
Wnt ligand secretion-inhibition hinders DTP^Diap^/Wnt^H^ population enrichment. **A-C)** Western blot analysis of active- (non-phosphorylated) β-catenin in MDA-MB-231, MDA-MB-468, and PDC-BRC-101 (shPLKO vs. shPORCN#1) cell lines treated with DOC or CAR for 96h. **D-F)** Flow cytometry analysis displaying % Annexin V⁺/DAPI⁺ cells of MDA-MB-231, MDA-MB-468, and PDC-BRC-101 (shPLKO vs. shPORCN#1) cell lines treated with DOC or CAR for 96h. Unpaired t-tests (n = 4 independent experiments). Data are presented as Mean ± SEM. **G)** Schematic representation of the Sequential Treatment (first IWP-2 pre-treatment followed by a recovery period then chemotherapy) model. **H)** Flow cytometry analysis displaying % of DTP**^Diap^**/Wnt**^H^** (GFP⁺) cells of MDA-MB-231-TGP, MDA-MB-468-TGP, and PDC-BRC-101-TGP cell lines pre-treated with IWP-2 (10uM) for 48h. **I)** Flow cytometry analysis of displaying % of DTP**^Diap^**/Wnt**^H^** (GFP⁺) cells of TNBC-TGP cell lines treated with DOC or CAR for 96h (with or without IWP-2 pre-treatment). Multiple t-tests corrected for multiple comparisons using the Holms-Sidak method (n = 3 independent experiments). Data are presented as Mean ± SEM. **J)** Schematic representation of the Combinatorial Treatment model. **K)** Flow cytometry analysis displaying % of DTP**^Diap^**/Wnt**^H^** (GFP⁺) cells of TNBC-TGP cell lines treated with DOC or CAR for 96h (sole or in combination with IWP-2). Unpaired t-tests (n = 4 independent experiments). Data are presented as Mean ± SEM. **L)** Flow cytometry analysis displaying % of Annexin V^+^ cells of TNBC cell lines treated with DOC or CAR for 96h (sole or in combination with IWP-2). Unpaired t-tests (n = 4 independent experiments). Data are presented as Mean ± SEM. **M)** Gene expression levels obtained via RT-qPCR of MYC for MDA-MB-231 and MDA-MB-468 cell lines treated with DOC or CAR for 96h (sole or in combination with IWP-2). Unpaired t-tests based on relative expression values (2^-dCt^) (n = 3 independent experiments). p values are indicated as *p < 0.05, **p < 0.01, ***p < 0.001, ****p < 0.0001, and ns, not significant.

Although several pharmacological inhibitors targeting the acyltransferase PORCN have been developed and shown efficacy in suppressing Wnt signaling in various solid tumors, notably in Wnt-deregulated colon cancers, the ongoing clinical trials investigating PORCN inhibitors in TNBC do not currently consider their interaction with chemotherapy^51–54^. Given our previous findings that the Wnt signaling pathway is activated in response to chemotherapeutic treatment, we sought to investigate whether pharmacological inhibition of PORCN could also prove effective in curbing Wnt-activation and induction of DTP**^Diap^/**Wnt**^H^**cells under treatment pressure. We examined two distinct approaches to therapeutically target treatment-induced DTP**^Diap^/**Wnt**^H^**cells. In the first approach, we pre-treated MDA-MB-231, MBD-MB-468, and PDC-BRC-101 TNBC cell lines with the Inhibitor of Wnt Production^55^ (IWP-2) for 48h followed by the application of either DOC or CAR agents (sequential treatment). In the second approach, we applied chemotherapeutic treatment simultaneously and in combination with IWP-2 (combinatorial treatment). Pre-treatment (sequential treatment strategy – Fig. 6G) of MDA-MB-231-TGP, MBD-MB-468-TGP, and PDC-BRC-101-TGP TNBC cell lines with IWP-2 for 48h led to a notable and significant reduction in the percentage of DTP**^Diap^/**Wnt**^H^** cells (Fig. 6H) with no effects on cell viability (Supplementary Fig. S6H). However, upon chemotherapy addition, we observed robust chemotherapy-induced Wnt-activation both in IWP-2 pre-treatment conditions and in sole chemo-treatment alike, seen in all TNBC cell lines (Fig. 6I). Notably, pre-treatment with IWP-2 followed by chemotherapy addition did not result in increased cell death nor sensitization to either agent (Supplementary Fig. S6I-K), suggesting that a sequential treatment strategy to target *a priori* existent DTP**^Diap^/**Wnt**^H^** population might not be sufficient to prevent chemotherapy-induced Wnt-activation and its subsequent implications.

In the second approach (combinatorial treatment strategy – Fig. 6J), simultaneous treatment of TNBC cell lines with either DOC or CAR therapeutic agent in combination with IWP-2 led to a significant decrease in the percentage of DTP**^Diap^/**Wnt**^H^** cells (Fig. 6K) underscoring the critical role of Wnt ligand-secretion in this acquired Wnt-activation phenomenon. Intriguingly, combinatorial treatment involving chemotherapeutic agents alongside IWP-2 resulted in a substantial increase in apoptotic cell death across all analyzed TNBC cell lines compared to treatment with either chemotherapeutic agent alone (Fig. 6L). Interestingly, we observed that the supplementation of IWP-2 alongside chemotherapy resulted in a significant rescue of MYC levels (Fig. 6M). Furthermore, using a second pharmacological PORCN inhibitor, WNT-974^53,54,56^ (LGK-974) resulted in similar outcomes obtained with Wnt ligand secretion-inhibitor IWP-2 (Supplementary Fig. S6L-Q).

In summary, our findings collectively demonstrate that Wnt ligand-secretion plays a crucial role in driving the enrichment of DTP**^Diap^/**Wnt**^H^**cells induced by chemotherapy. Notably, simultaneous-combinatorial treatment, rather than sequential, encompassing chemotherapeutic agents and pharmacological inhibitors of Wnt ligand-secretion can significantly reduce the induction and/or enrichment of DTP**^Diap^/**Wnt**^H^** cells and enhance the sensitivity of TNBC cell lines to chemotherapy.

### Inhibition of Wnt ligand-secretion and chemotherapeutic treatment synergistically sensitize *in vivo* xenograft TNBC model to treatment

A precise and complete understanding of Wnt-activation kinetics and dynamics in response to chemotherapeutic treatment in *in vivo* models is lacking. To shed some light on this phenomenon, we engineered the TNBC cell line MDA-MB-231 with the Wnt-transcriptional reporter TOPFLASH^35^, a β-catenin-responsive firefly luciferase reporter plasmid compatible for use with the *in vivo* live imaging system (IVIS) (Fig. 7A).

**Fig. 7:**
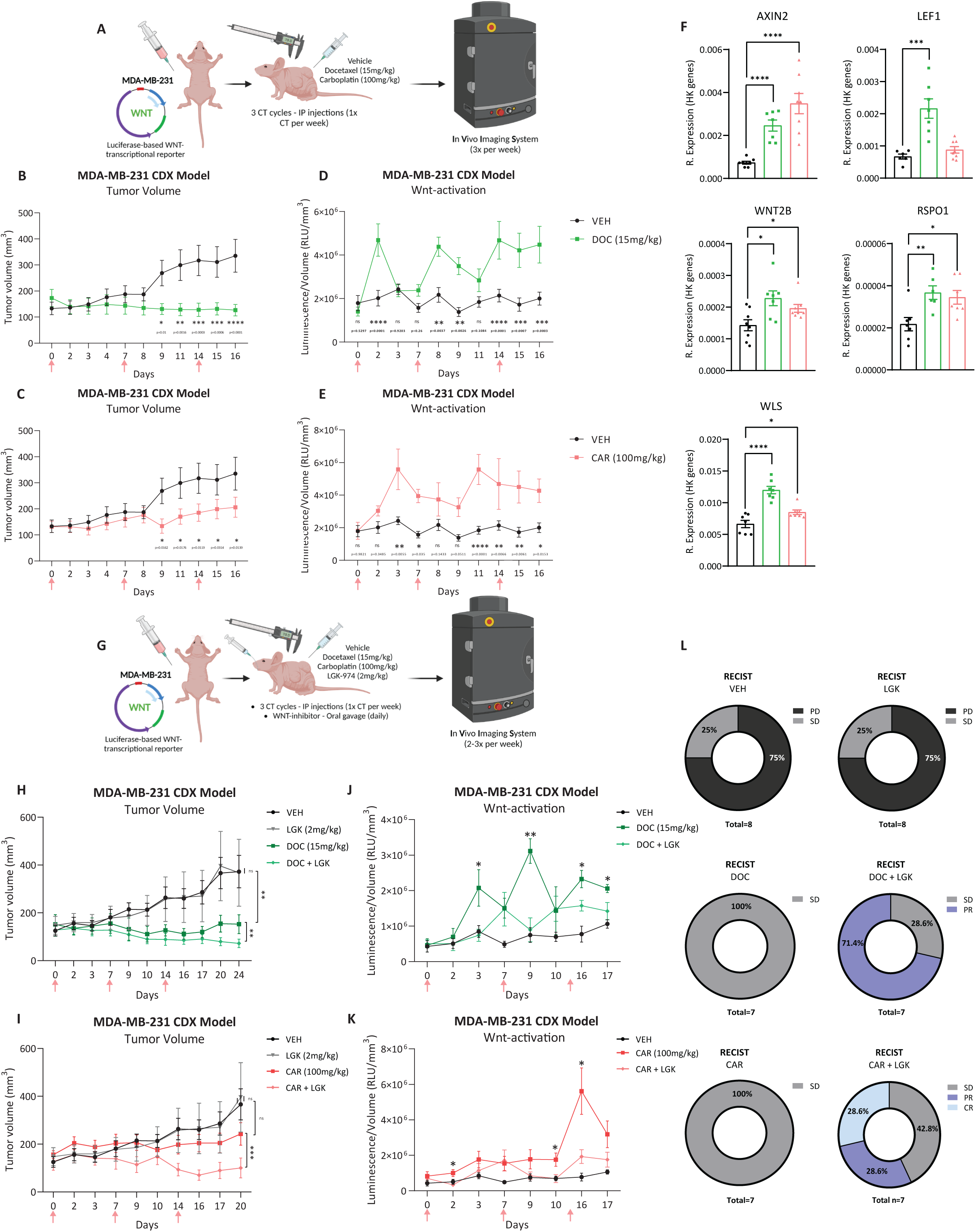
Inhibition of Wnt ligand-secretion and chemotherapeutic treatment synergistically sensitize *in vivo* xenograft TNBC model to treatment. **A)** Schematic representation of cell line derived xenograft (CDX) experimental setup to study Wnt signaling pathway kinetics upon chemotherapeutic treatment *in vivo*. **B-C)** Tumor growth curves of subcutaneous generated xenograft models treated with VEH, DOC (15mg/kg/week – top), or CAR (100mg/kg/week – bottom). Pink arrows indicate administration of chemotherapy. Two-way ANOVA with Fisher’s LSD test (n = 7 mice for all treatment groups). Data are presented as Mean ± SEM. **D-E)** Levels of Wnt-activation (RLU/mm3) displayed as luminescent signals (RLU) captured by IVIS Spectrum normalized to tumor volume (mm3) in xenograft models treated with VEH, DOC (top), or CAR (bottom). Two-way ANOVA with Fisher’s LSD test (n = 7 mice for all treatment groups). Data are presented as Mean ± SEM. **F)** Gene expression levels obtained via RT-qPCR of various Wnt-targets and Wnt-activators in samples resected from xenograft models treated with VEH, DOC, or CAR. Unpaired t-tests based on relative expression values (2^-dCt^) (n = 7 mice for all treatment groups). Data are presented as Mean ± SEM. **G)** Schematic representation of CDX experimental setup with Wnt ligand secretion inhibition *in vivo*. **H-I)** Tumor growth curves of xenograft models treated with VEH, LGK (2mg/kg/day), DOC (top), DOC+LGK (top), CAR (bottom), and CAR+LGK (bottom). Pink arrows indicate administration of chemotherapy. Paired t-tests (based on final tumor volumes – obtained on day of sacrifice, n = 8,7,6,5 mice per treatment group). Data are presented as Mean ± SEM. **J-K)** Levels of Wnt-activation (RLU/mm3) displayed as luminescent signals (RLU) captured by IVIS Spectrum normalized to tumor volume (mm3) in xenograft models treated with VEH, DOC (top), DOC+LGK (top), CAR (bottom), and CAR+LGK (bottom). Multiple t-tests (n = 8,7,6,5 mice per treatment group). Data are presented as Mean ± SEM. **L)** RECIST analysis displayed as percentage of animals per treatment group (VEH, LGK, DOC, DOC+LGK, CAR, and CAR+LGK) classified/assigned to one of four distinct tumor responses to treatment: Progressive Disease (PD), Stable Disease (SD), Partial Response (PR), or Complete Response (CR). p values are indicated as *p < 0.05, **p < 0.01, ***p < 0.001, ****p < 0.0001, and ns, not significant.

Treatment with either DOC or CAR^57,58^ resulted in a significant overall decrease in tumor volume when compared to the vehicle (VEH) treated group (Fig. 7B, C). Notably, at the administered doses, no significant changes in mouse body weight were observed during the three weeks of treatment (Supplementary Fig. S7A). We observed, as early as 48h and 72h (for DOC and CAR, respectively) upon chemotherapeutic administration, a significant upregulation in transcriptional Wnt-activation recorded by IVIS (Fig. 7D, E, and representative image Supplementary Fig. S7B). Notably, this activation started to decrease as the week progressed following the 1^st^ dose-administration and again increased significantly 48h and 72h (for DOC and CAR, respectively) following administration of the 2^nd^ and 3^rd^ dose (Fig. 7D, E) highlighting the dynamic behavior of chemotherapy-induced Wnt-activation *in vivo*. Gene expression analysis on the resected tumors following the 3^rd^ and final cycle of chemotherapeutic administration revealed elevated expression of Wnt-targets (*AXIN2* and *LEF1*), Wnt ligands and -enhancers (*WNT2B*, *WNT3A*, *WNT7B*, *RSPO1*, and *RSPO3*), and Wnt secretion machinery components (*WLS* and *PORCN*) (Fig. 7F and Supplementary Fig. S7C) in chemotherapy-treated groups.

We next proceeded to assess whether a combinatorial treatment strategy encompassing the use of chemotherapy with a pharmacological Wnt ligand secretion-inhibitor (Fig. 7G) could abrogate Wnt-activation and lead to tumor sensitization, as indicated in our previous *in vitro* findings (Fig. 6K, L and Supplementary Fig. S6L-Q). Administration of LGK-974 alone had no discernible effect on tumor growth^56,59^ (Fig. 7H, I, and representative images Supplementary Fig. S7D). Conversely, the combination of LGK-974 with either DOC or CAR treatment resulted in a substantial and significant decrease in Wnt pathway-activation, correlating with a marked reduction in tumor volume compared to solo chemo-treated or VEH-groups (Fig. 7H-K and representative images Supplementary Fig. S7D). Notably, LGK-treatment, sole or in combination with either chemotherapeutic agent, had no significant effect on mouse body weight observed during the three weeks of treatment (Supplementary Fig. S7E).

Response Evaluation Criteria in Solid Tumors^60^ (RECIST) analysis was performed to assess tumor response to treatment, categorizing objective outcomes into progressive disease (PD), stable disease (SD), partial response (PR), and complete response (CR). In the VEH-group, 75% (6/8) of tumors were classified as PD, and 25% (2/8) were classified as SD (Fig. 7L). Sole LGK-974 treatment showed no difference in RECIST classifications (75% PD and 25% SD) compared to VEH-conditions, indicating minimal impact on tumor burden (Fig. 7L). In solo DOC- or CAR-treated groups, RECIST analysis classified 100% of tumors as SD, demonstrating the efficacy of DOC or CAR treatments in controlling tumor growth (Fig. 7L). In the DOC+LGK treatment arm, LGK-974 supplementation significantly improved objective response with tumors classified as 28.6% SD (2/7) and 71.4% PR (5/7) (Fig. 7l), compared to sole DOC-treatment (100% SD). Similarly, in the CAR+LGK treatment arm, tumors were classified as 42.8% SD (3/7), 28.6% PR (2/7), and 28.6% CR (2/7), highlighting the positive impact of combining chemotherapy with Wnt secretion-inhibition (Fig. 7L).

Our study comprehensively characterizes the activation dynamics of the Wnt signaling pathway in chemotherapy-treated tumors within an *in vivo* setting. We demonstrate that activation of the Wnt pathway is primarily triggered by Wnt ligand-secretion as a combined treatment approach, which includes chemotherapy and Wnt ligand secretion-inhibition, significantly reduces Wnt pathway-activation and effectively curbs tumor growth.

### Preclinical PDO models recapitulate chemotherapy-mediated Wnt-activation and sensitization to synergistic Wnt ligand secretion-inhibition

Transcriptomic analysis performed on RNA-sequencing-based datasets of longitudinally paired samples of breast cancers patients during NAC treatment^61^ (GSE123845 – Fig. 8A and Supplementary Table 6) showed that an in-house derived (Wnt^High^) Wnt-signaling signature was significantly enriched in tumor samples obtained from patients undergoing (on-NAC) NAC treatment (Fig. 8B). Parallelly, and in conjunction with our findings in this study using *in vitro* cell lines, enrichment of Wnt-signaling was seen hand in hand with an enrichment of the Rehman et. al. DTP**^Diap^** gene signature and a suppressed MYC hallmark signature (Fig. 8C, D), further highlighting the interplay between the Wnt^High^ and DTP**^Diap^** phenotypes in a clinical setting. Next, we investigated the effects of chemotherapeutic treatment on the Wnt-signaling pathway in pre-clinical 3D PDO models^62,63^. Two different PDO models, R1-IDC113 (113 BCO) and R2-IDC159A (159A BCO) (Representative images Fig. 8E and Supplementary Fig. S8A) were used. Typically, organoid models are cultured in growth factor-rich medium^64^, of which Wnt ligand (Wnt-3a) and Wnt ligand-enhancer (R-spondin3) are components, possibly influencing studies of Wnt signaling pathway dynamics. Culturing of either cancer organoid model for four passages in a Wnt^-^/Rspo^-^ breast cancer organoid (BCO) medium, had no effects on the morphology, proliferation rate, or viability of either PDO model when compared to a baseline (Wnt^+^/Rspo^+^) BCO medium^65^ (Supplementary Fig. S8B, C). Upon exposure to IC50 concentrations of either DOC or CAR agents (Supplementary Fig. S8D-G) in a Wnt^-^/Rspo^-^ BCO medium, both PDO models exhibited a substantial and statistically significant increase in the expression levels of Wnt-target genes, compared to UNT conditions (Fig. 8F, G). Immunofluorescence analysis of active β-catenin levels confirmed increased Wnt-activation in both PDO models following exposure to either chemotherapeutic agent (Fig. 8H and Supplementary Fig. S8H).

**Fig. 8:**
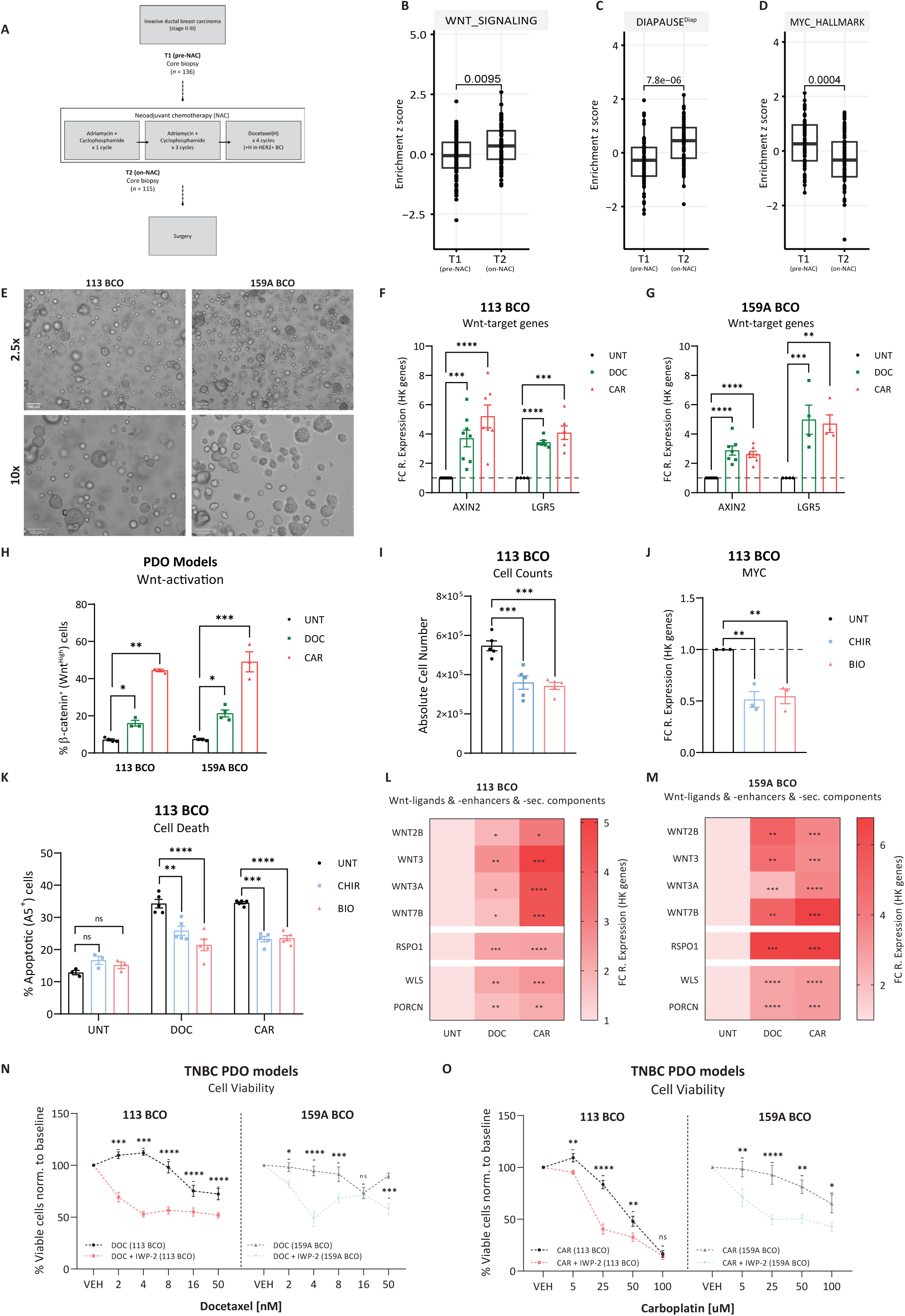
Preclinical PDO models recapitulate chemotherapy-mediated Wnt-activation and sensitization to synergistic Wnt ligand secretion-inhibition. **A)** Overview of supplementary information of patient dataset (GSE123845) analyzed in this study. **B-D)** Association analyses and enrichment scores of an in-house derived Wnt^High^ signature, DTP**^Diap^** signature^7^, and MYC GSEA hallmark signature evaluted in core biopsies pre-NAC and during (on)-NAC. **E)** Phase-contrast images of TNBC-PDO models, 113 BCO (left) and 159A BCO (right) in basal culture conditions at 2.5x (top) and 10x (bottom) magnification **F-G)** Gene expression levels obtained via RT-qPCR of Wnt-targets (AXIN2 and LGR5) for 113 BCO and 159A BCO models treated with DOC (16nM for 113 BCO and 8nM for 159A BCO) or CAR (50µM for 113 BCO and 125µM for 159A BCO) for 96h, displayed as fold change (to UNT) of 2^-dCt^ values (relative to HK-genes). Multiple t-tests corrected for multiple comparisons using the Holm-Sidak method based on relative expression values (2^-dCt^) (n = 4 independent experiments). **H)** Flow cytometry analysis displaying % of Wnt-active (β-catenin⁺) cells from 113 BCO (left) and 159A BCO (right) models treated with DOC or CAR for 96h. Multiple t-tests corrected for multiple comparisons using the Holm-Sidak method (n = 3 independent experiments). Data are presented as Mean ± SEM. **I)** Absolute cell number of 113 BCO model treated with CHIR or BIO for 72h. Unpaired t-tests (n = 3 independent experiments). Data are represented as Mean ± SEM. **J)** Gene expression levels obtained via RT-qPCR of MYC for 113 BCO model treated with CHIR or BIO for 72h, displayed as fold change (to UNT) of 2^-dCt^ values (relative to HK-genes). Unpaired t-tests (n=3 independent experiments). Data are represented as Mean ± SEM. **K)** Flow cytometry analysis displaying % of Annexin V⁺ cells of 113 BCO model treated DOC or CAR for 96h (sole or pre-treated with CHIR or BIO for 48h). Two-way ANOVA corrected for multiple comparisons using Tukey’s test (n = 3 independent experiments). Data are represented as Mean ± SEM. **L-M)** Heatmaps showing gene expression levels obtained via RT-qPCR of Wnt-ligands, -enhancers, and -secretion machinery components for 113 BCO (left) and 159A BCO (right) models treated with DOC or CAR for 96h, displayed as fold change (to UNT) of 2^-dCt^ values (relative to HK-genes). Unpaired t-tests based on relative expression values (2^-dCt^) (n = 4 independent experiments). **N-O)** Drug dose-response curves of 113 BCO and 159A BCO models treated with increasing concentrations of DOC (right) or CAR (left) (sole or in combination with IWP-2, 50uM). Viability in sole- or combinatorial-treatment is normalized to UNT or sole-IWP-2 conditions (baseline). Multiple t-tests corrected for multiple comparisons using the Holm-Sidak method (n = 4 independent experiments). Data are presented as Mean ± SEM. p values are indicated as *p < 0.05, **p < 0.01, ***p < 0.001, ****p < 0.0001, and ns, not significant.

To evaluate if activation of the Wnt signaling pathway would also suffice in inducing a growth-arrest, drug-tolerant, DTP-state in a pre-clinical TNBC PDO model, we treated the 113 BCO PDO model with CHIR or BIO to stimulate activation of the Wnt signaling pathway as previously done using TNBC cell lines. Treatment with either CHIR or BIO resulted in transcriptional activation of the Wnt signaling pathway as evidenced by a significant increase in Wnt-target gene expression (*AXIN2* – Supplementary Fig. S8I). Parallelly, and in accordance with our previous findings, treatment with CHIR or BIO led to a significant decrease in cell number as a possible consequence of growth-arrest (Fig. 8I) seen in conjunction with significant suppression of MYC activity (Fig. 8J and Supplementary Fig. S8J). Furthermore, pre-treatment with either CHIR or BIO was able to induce a drug-tolerant phenotype in the 113 BCO PDO model, whereby we observed a significant decrease in apoptosis induction in cells pre-treated with CHIR or BIO followed by chemotherapeutic treatment when compared to sole chemotherapy exposure (Fig. 8K).

Next, we confirmed significant elevation in the expression levels of Wnt ligands, -enhancers, and -secretion machinery components in chemotherapy-treated PDO models when compared to UNT conditions (Fig. 8L, M). To investigate the efficacy of the combinatorial treatment strategy, both PDO models were exposed to Wnt ligand secretion-inhibition alone and in combination with escalating concentrations of either chemotherapeutic agent for 96h. Treatment of both PDO models with IWP-2 alone for 96h did not have any effect on cell viability or proliferation (Supplementary Fig. S8K). However, the exposure of both PDO models to chemotherapy in combination with IWP-2 demonstrated a significant reduction in the percentage of viable cells compared to chemotherapeutic treatment alone (Fig. 8N, O). Interestingly, this sensitization effect was most effective when IWP-2 was supplemented with sublethal concentrations of chemotherapy (<16nM DOC and <50µM CAR – 113 BCO | <8nM DOC and <125µM CAR – 159A BCO).

In summary, our study demonstrates chemotherapy-induced Wnt-activation in TNBC pre-clinical PDO models and in transcriptomic datasets derived from patients samples undergoing NAC. Notably, TNBC PDO models exhibited a robust and enhanced sensitization to the combinatorial treatment comprising Wnt ligand secretion-inhibition alongside sub-lethal chemotherapy (<determined IC50) concentrations. This finding underscores the potential clinical significance of this combinatorial approach for breast cancer treatment.

## DISCUSSION

Understanding the introductory events that lead to the formation of DTP**^Diap^** cells could provide new therapeutic strategies to prevent the development of drug-resistant populations even before they are steadily established.

Our findings reveal a significant enrichment in the percentage of a Wnt^High^ population alongside an increase in the intensity of Wnt-activation in various TNBC models subjected to distinct chemotherapeutic agents. We establish a significant transcriptional association between the Wnt^High^ population and DTP**^Diap^** population, including a unique DTP**^Diap^** gene-signature and a reduction of MYC-targets hallmark; both recently correlated with cancer DTP cells^7,11^. Functional analyses confirmed that Wnt^High^ cells exhibit DTP**^Diap^** cell-traits such as reduced proliferation and an enhanced capacity for drug tolerance, indicating that Wnt^High^ cells function as *bona fide* DTP**^Diap^** cells. We find that the transcriptional activation of the Wnt signaling pathway serves as a distinctive biomarker for the early emergence of the DTP**^Diap^** cell-phenotype in untreated parental cells, but particularly in cells responding to chemotherapeutic exposure.

Enrichment of DTP cells has been observed across various chemotherapeutic treatments. In our study, we utilize two chemotherapeutics with distinct mechanisms of action: Docetaxel, which induces microtubule stabilization leading to cell cycle arrest and apoptosis, and Carboplatin, which forms DNA crosslinks that inhibit DNA replication and transcription, ultimately leading to cell death. Despite their different pro-apoptotic mechanisms, both treatments result in an enrichment of DTP**^Diap^/**Wnt**^H^** cells, showing that distinct chemotherapies converge on the enrichment of Wnt ligand-expression and activation of the Wnt signaling pathway. Interestingly, in regenerative models, such as Hydra, increased Wnt ligand-expression has been noted in cells undergoing apoptosis as a pro-survival mechanism in response to tissue damage^66^. This suggests that, under the stress of chemotherapeutic pressure, cells may activate similar pro-survival mechanisms and signaling pathways. In this article, we propose a mechanistic model for acquired chemoresistance in TNBC mediated by the enrichment of drug-tolerant cells, involving two phases. Initially, cells sense environmental changes induced by therapeutic pressure, leading to elevated levels of Wnt ligands and -enhancers, along with increased Wnt ligand-secretion. Subsequently, cells adapt to these pressures by transcriptionally activating the Wnt signaling pathway, ultimately resulting in the enrichment of DTP**^Diap^/**Wnt**^H^**cells.

DTP**^Diap^** cells have been found to be susceptible to ferroptosis induction, a form of programmed cell death dependent on iron and characterized by the accumulation of lipid peroxides^67^. Recently, BET inhibitors have also emerged as a promising therapeutic strategy for targeting and eliminating persister cells^68^. However, current research predominantly focuses on identifying and exploiting vulnerabilities in existing DTP cells or in drug-tolerant expanded persisters (DTEPs) that have resumed cell division after a period of drug withdrawal. Consequently, these therapeutic agents are designed to target DTP**^Diap^** cells specifically, rather than preventing the acquisition and emergence of a DTP**^Diap^** cell-state. Our results show that combinatorial treatment involving Wnt ligand secretion-inhibition supplemented alongside chemotherapy reduces DTP**^Diap^/**Wnt**^H^** cell-enrichment and sensitizes tumors to treatment, holding significant promise for future clinical trials in TNBC. Surprisingly, current clinical trials investigating PORCN inhibition in TNBC are restricted only to Wnt-deregulated cancers and exclude consideration of chemotherapy (NCT03447470 and NCT01351103)^53,54^. Our study also underscores the importance of temporal considerations in treatment regimens. Crucially, we show that pre-treatment with PORCN inhibitors does not prevent a substantial increase in DTP**^Diap^/**Wnt**^H^** cells once chemo-treatment is applied, indicating that patients undergoing chemotherapy might not benefit from an initial treatment with Wnt-inhibitors. Therefore, our findings suggest that simultaneous-combinatorial treatment, rather than sequential treatment, encompassing Wnt-inhibitors and chemotherapy might provide a solution to effectively control and prevent chemotherapy-induced drug tolerant cell-enrichment while simultaneously sensitizing tumors to the effects of chemotherapy.

The origin of DTP cells is a subject of ongoing debate. Some research suggests that DTP cells may have a stable clonal origin, while others propose that DTP cells enter a temporary drug-tolerant state under the influence of external factors such as chemotherapy. Our live-cell imaging studies, which allow the tracking and tracing of Wnt-reporter TNBC cell lines, indicate that chemotherapy treatment leads to a significant enrichment of DTP**^Diap^/**Wnt**^H^** cells, primarily in cells that were initially in a Wnt^Low^ state. This suggests a vital role of *de novo* activation of the Wnt signaling pathway in response to chemotherapeutic treatment. Additionally, a lower but notable proportion of DTP**^Diap^/**Wnt**^H^**cells in chemo-treated conditions were initially in a transcriptionally Wnt-active state. These observations imply that both intrinsic and acquired resistance mechanisms, driven by Wnt transcriptional-activity, coexist and contribute to DTP cell-formation. Importantly, our results show that activation of the Wnt siangling pathway through GSK3 inhibition is sufficient to induce a DTP**^Diap^** state in parental breast cancer cells, even in the absence of therapeutic pressure. This demonstrates that the DTP**^Diap^** cell-state is not exclusive to chemotherapy-treated samples and can emerge under non-challenging conditions, explaining its presence in untreated populations.

Activation of the canonical Wnt signaling pathway and β-catenin stabilization have been correlated with unfavorable prognosis in patients with TNBC^47,69–73^. Wnt-activation has been extensively associated with induction of proliferation, increased MYC expression^74,75^ and a CSC phenotype^56,69,70,73,76^. Contrary to previous research, our results establish a significant transcriptional link between the Wnt^High^ population and the slow proliferating DTP**^Diap^** cells with reduced MYC expression. The reductive proliferation effect of Wnt-activation in TNBC is reminiscent of the effect of LGR5^+^ cells in mouse models of basal cell carcinoma (BCC), in which therapy treated LGR5^+^ BCC cells correlate with increased Wnt-activity and a slow-proliferative phenotype^77^, suggesting that, in other cancer types, Wnt-activation might also promote a diapause-like state.

Prior studies have shown that Wnt-activation reduces the proliferation of mouse pluripotent ESCs by increasing the expression of cyclin-dependent kinase inhibitors (CDKis) such as p16^Ink4a^, p21, and p27, alongside a decrease in MYC expression^78,79^. Whether Wnt-activation might also induce a diapause state in mESCs needs further investigation, however RNA sequencing of dormant versus reactivated mouse blastocysts identified differential regulation of the Wnt signaling pathway^80^. This finding is consistent with the work of Fan et al., who showed higher Wnt transcriptional-activity in epiblast cells of hormonally diapaused embryos compared to actively cycling embryos^81^. This suggests that the Wnt signaling pathway may play a direct role in promoting a diapause cell state also in preimplantation embryos or pluripotent cells.

In summary, our research suggests that Wnt pathway-activation plays a pivotal role as an early event enhancing the enrichment of DTP**^Diap^/**Wnt**^H^** cell populations in TNBC and contributing to the development of therapeutic resistance, particularly in response to chemotherapy. Therefore, a potential strategy to address DTP**^Diap^/**Wnt**^H^** cell enrichment could involve targeting Wnt ligand-secretion successively hindering chemotherapy-induced Wnt/β-catenin pathway-activation and subsequently diminishing the sustenance and enrichment of DTP cell-populations.

## Supporting information

Supplementary Table 1 - DESEQ

Supplementary Table 2 - GSEA

Supplementary Table 3 - DESEQ

Supplementary Table 4 - Regression model (Forrest Plots)

Supplementary Table 5 - Wilcox Plots

Supplementary Table 6 - GSE123845

## AUTHOR CONTRIBUTIONS

YEL, WO, and FL conceived and designed the study. WO partially carried out flow cytometry experiments, RT-qPCR experiments, analysis of publicly available datasets. AP performed all bioinformatic and biostatistical analyses under the supervision of FR and CD. AQ partially carried out flow cytometry experiments, western blot analysis, RT-qPCR experiments, and *in vivo* experiments. FR provided critical input for the bulk transcriptomic analysis under supervision of and in consultation with CD. PA participated in library preparations for mRNA-seq samples and provided input in experimental design. CL and WDW provided aid in the *in vivo* experiments under supervision of SS and DA. LM provided aid for the culturing and maintenance of patient derived organoid (PDO) models. SH provided input and feedback into experimental design under supervision of AB. SM generated the *in vitro* cell line, PDC-BRC-101. AdJS provided critical input in experimental design and assisted in the generation of several fluorescent reporter cell lines. MB provided input in the *in vivo* experimental design. SJS provided critical input in human patient dataset analysis. CS provided the PDO models. AB provided critical input for experimental design. DA provided critical input in *in vivo* data analysis. YEL carried out the remainder of the experimental work. Data analysis and figure preparation were performed by YEL and WO and reviewed by FL. The manuscript was written by YEL and FL and reviewed and approved by all authors. FL secured funding and supervised and guided experimental work and manuscript preparation.

## ACKNOWLEDGEMENTS

We are grateful to the KU Leuven FACS core team for providing the facility. We also thank the KU Leuven Genomics Core (http://genomicscore.be) for RNA sequencing, and data processing. We also thank the KU Leuven TRACE core and the MOSAIC core for help in *in vivo* experimental design and animal imaging. The authors would like to extend their gratitude to the FWO Research Foundation – Flanders for the Ph.D. fellowships awarded to P.A. (11M7822N). AdJS was supported by PhD Emmanuel van der Schueren (EvdS) fellowship granted by Kom Op Tegen Kanker (KOTK) and a Postdoctoral Mandate (PDM) granted by KU Leuven. The Lluis Lab is financed by the FWO Research Project Grants G091521N (AB, DA, FLL), G073622N (FLL), and the C1 KU Leuven internal grant C14/21/115 (FLL).

## MATERIALS AND METHODS

### Ethics declaration

All xenograft animal experiments performed were approved by the Ethics Committee at KU Leuven University under the ethical approval codes P055/2022 and P016/2023.

Patient derived organoid (PDO) models used in this study were established from freshly resected tumor tissues obtained from TNBC patients at the Antoni van Leeuwenhoek Hospital. The study was approved by the institutional review board (NKI-B17PRE) and the subjects provided informed consent.

All cell lines used in this study are approved for use by the Ethics Committee at the KU Leuven University Biobank under the code S65166.

### TNBC cell line culture

MDA-MB-231 (ATCC-HTB-26) and MDA-MB-468 (ATCC-HTB-132) were maintained in DMEM high glucose (Gibco, 41965039) supplemented with 10% (v/v) fetal bovine serum, 1mM sodium pyruvate (Gibco, 11140035), 100µg/mL penicillin-streptomycin (Gibco, 15140163), and 0.01mM 2-mercapthoethanol (Gibco, 31350010).

PDC-BRC-101 cell line (PDX-derived cell line) was obtained from collaborators, Daniela Anibali and Stijn Moens (Amant Lab – Gynecological Oncology) – KU Leuven and maintained in OCMI media^82^ composed of a composed of 1:1 mixture of Medium199 (Gibco, 31150022) and DMEM F-12 (Gibco, 11320074) supplemented with 10% (v/v) fetal bovine serum, 100µg/mL penicillin-streptomycin, 20µg/mL insulin (Sigma/Merck, I9278), 25ng/mL cholera toxin subunit B (Sigma/Merck, C9903-.5MG), 0.5µg/mL hydrocortisone (Sigma/Merck, H0888-1G), and 10ng/mL epidermal growth factor (Stem Cell Technologies, 78006.1).

All cell lines were cultured in 84mm x 20mm (D x H) tissue-culture treated dishes at 37°C and 5% CO_2_ and maintained at 70-80% confluency. For cell line passaging and plating, 1X phosphate-buffered saline (Gibco, 10010-015) was used as a washing solution followed by dissociation using 0.25% trypsin-EDTA (Gibco, 25200-056) and cell-pelleting by centrifugation for 4 minutes at 300G (0.3rcf). Cells were counted manually via the BRAND counting chamber Neubauer improved (Sigma Aldrich/Merck, BR717810-1EA) under a 10X objective lens using a Leica DMi inverted microscope. The same microscope, equipped with a 2.5 Megapixel HD Microscope Camera Leica MC120 HC, was used to obtain images of cultured cancer cell lines. Unless specified otherwise, cancer cell lines were plated according to the following seeding densities: 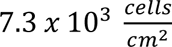 and 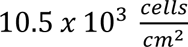 (MDA-MB-231 and MDA-MB-468/PDC-BRC-101, respectively).

### Chemotherapeutic and small molecule treatment of TNBC cell lines and PDO models

Cell lines were treated with increasing concentrations of docetaxel (Taxotere, 0–144 nM) and carboplatin (Carbosin, 0–1600 µM) for 72h. Cell metabolic activity, reflecting cell-number and -viability) was assessed using the MTT [Thiazolyl Blue Tetrazolium Bromide] assay (Sigma/Merck, M5655-500mg) according to manufacturer’s instructions and sigmoidal dose-response curves were generated to calculate the mean IC50 values of each drug that were used in the subsequent study. Chemotherapeutic agents were obtained from the pharmacy of Universitair Ziekenhuis (UZ) Leuven.

For Wnt pathway-stimulation, CHIR99021 (CHIR – Sigma/Merck, SML1046) and 6-Bromoindirubin-3’-oxime (BIO – Sigma/Merck, B1686-5MG) were used at 8µM and 3µM, for CHIR and BIO, respectively.

### Lentiviral particle production and transduction

Lentiviruses were produced according to the RNAi Consortium (TRC) protocol available from the Broad Institute (https://portals.broadinstitute.org/gpp/public/resources/protocols). In brief, 7 × 10^5^ HEK-293T cells were seeded per well in 6-well plates and transfected the following day with 750 µg pCMV-dR8.91, 250 µg pCMV-VSV-G, and 1 µg of the specific lentiviral plasmid/construct using FugeneHD (Promega, E2311) in Optimem (Gibco, 31985070). One day after, the culture medium was refreshed. The same day, lentivirus-recipient cells were plated in 6-well plates at their respective concentrations (see cell line culture). Lentivirus-containing medium was collected from HEK293T cells 48h and 72h post-transfection and added to recipient cancer cells after filtration using a 0.45 µM filter (VWR-Corning, 431220). 48h post infection, recipient cancer cells were washed thoroughly with PBS, medium refreshed, and the appropriate selection antibiotics applied until selection process was completed.

Wnt-transcriptional reporters, TOPGFP (7xTcf-eGFP // SV40-PuroR), TOPFLASH (7xTcf-FFluc), and mCherry-TOPGFP (7xTcf-eGFP // SV40-mCherry) were obtained from Addgene (#24305, #24308, and #24304, respectively). Wnt-transcriptional reporter dTOPGFP (dTGP) was gifted to us from the Moon lab University of (Washington – USA).

For PORCN shRNA mediated silencing, we used the MISSION® Lentiviral shRNA (Sigma Aldrich/Merck, SHCLNG – clones, TCRN000153848 and TCRN000157366) and the MISSION pLKO.1-puro Non-Target shRNA Control Plasmid DNA (SHC016-1EA) as a negative control in experiments.

### Real-Time Quantitative Polymerase Chain Reaction and Gene Expression Analysis

For RT-qPCR, total RNA was extracted (from TNBC cell lines or cryopreserved tumor tissue) using the GenElute mammalian total RNA miniprep kit (Sigma/Merck, RTN350-1KT) according to manufacturer’s instructions with an additional step of DNA digestion using the On-Column DNAse I digestion set according to manufacturer’s instructions (Sigma/Merck, DNASE70). cDNA was synthesized from 500 ng of total RNA using the BIORAD iScript cDNA cDNA synthesis kit (BIORAD, CAT#1708891), according to manufacturer’s instructions. Quantitative real-time PCR reactions were set up in technical triplicates with Platinum SYBR Green qPCR SuperMix-UDG (Invitrogen, 11733-046) on a ViiA7 Real-Time PCR System (Thermo-Scientific). Expression levels were normalized to two housekeeping genes (HK) RPL19 and GAPDH to determine ΔCT values. Statistical testing of differences in expression levels between samples was carried out based on relative-expression values (2^−Δ*CT*^). In some figures, gene expression values are represented as fold-change for convenience of interpretation, although statistical testing was performed on relative expression values (2^−Δ*CT*^).

### SDS-PAGE and Western Blot analysis

TNBC cell lines were washed with PBS and collected/pelleted by centrifugation. Whole cell lysates were obtained via mechanical lysis using a needle (VWR-TERUMO, AN2138R1) and RIPA cell lysis buffer (Sigma/Merck, R0278-50mL) supplemented with a cocktail of 1:100 phosphatase inhibitors cocktail 2 and 3 (Sigma/Merck, P5726-1ML and P0044-1ML, respectively) and 1:100 protease inhibitor cocktail (Sigma/Merck, 11873580001). Samples were placed on a rotation wheel for a minimum of 30 minutes at 4 °C after which they were centrifuged at 16,000x g for 10 minutes at 4 °C. The supernatant from the lysates was collected and protein concentration was determined using the Bradford Assay (Biorad, 5000006). For SDS-PAGE 20 mg of protein were mixed with 4x Laemmli buffer (240 mM Tris/HCL pH 6.8, 8% SDS, 0.04% bromophenol blue, 5% 2-mercaptoethanol, 40% glycerol) and denatured for 5 minutes at 95°C prior to electrophoretic protein separation. Resolved protein extracts were transferred to PVDF membranes (BIORAD, 162-0177). Transfer success was assessed with Ponceau S solution, and membranes were blocked with 5% non-fat milk or 5% BSA in TBS-T (0.1% Tween-20®) for 60 minutes. After blocking, membranes were incubated with primary antibodies at 4°C overnight. The day after, membranes were washed 3 times with PBS-T for 10 minutes and incubated with secondary HRP-conjugated antibodies. Immunolabeled proteins were detected with Supersignal West Pico chemiluminescent kit (Fisher Scientific, 34077) on autoradiography film (Santa Cruz, SC-201697). The primary antibodies used were active rabbit anti-non-phosphorylated β-catenin (CellSignaling Technologies, #19807S), rabbit anti-PORCN (Novus Biologicals, NBP1-59677), and rabbit anti-WNT2b (Abcam, ab178418). Mouse anti-β-Actin (Santa Cruz Biotechnology; sc-47778) was used as a loading control.

### Flow Cytometry

For Wnt-activation assessment, cells were washed with PBS and collected/pelleted by centrifugation. Cells were resuspended in PBS2%FBS, counterstained with 5µg of 4’,6-diamidino-2-phenylindole (DAPI – 1:1) (Sigma/Merck, D9542-10mg) to eliminate dead cells before running through the flow cytometer. Cell lines lacking any of the previously described Wnt-transcriptional reporters were used as gating controls.

For ALDH activity assay, cells were washed with PBS and collected/pelleted by centrifugation. Cells were stained using the AldeRed ALDH detection assay (Sigma/Merck, SRC150) according to manufacturer instructions. Cells were counterstained with 5µg of DAPI (1:1) to eliminate dead cells before running through the flow cytometer.

For immunolabeling of CD44 and CD24, cells were detached, washed twice in PBS with 4% FBS, and incubated with CD44-PE (BD Pharmigen, 555479) and CD24-APC (Invitrogen, 17-4714-81) antibodies according to manufacturer specifications at room temperature. After incubations, cells were washed twice in PBS with FBS and resuspended in PBS containing 4% FBS and 100 nM of DAPI. Cells incubated with PE- and APC-conjugated isotype-antibodies and single-stained cells were used as gating controls.

For Annexin V apoptosis analysis, cells were washed with PBS and collected/pelleted by centrifugation. Cells were resuspended in 1x Annexin V binding buffer (BD Pharmigen, 51-66121E) and incubated at room temperature in the dark for 15 minutes with APC-conjugated Annexin V (Thermo-eBioscience, BMS306APC-100). After incubation, cells were diluted in 1X binding buffer supplemented with 100 nM of DAPI before running through the flow cytometer. Unstained and single-stained (Annexin V-only or DAPI-only stained) cells were used as gating controls.

To obtain chemotherapy-induced Wnt^High^ and Wnt^Low^ cells, cells were washed with PBS and collected/pelleted by centrifugation. Cells were resuspended in PBS4%FBS, counterstained with 5µg of DAPI (1:1) to eliminate dead cells before running through the SONY MA900 Multi-Application Cell Sorter. Depending on the application, 2 − 3 × 10^5^cells were sorted (based on their GFP expression) into 1.5mL Eppendorf tubs (with 300µl of PBS4%FBS) and either used for RNA-extraction and gene expression analysis or for re-culturing.

For immunostaining of active- (non-phosphorylated) β-catenin, cells were washed with PBS and collected/pelleted by centrifugation. Cells were fixed with ice-cold 70% Ethanol. After which samples were with PBS2%FBS and blocked with 5% donkey serum (in PBS) at room temperature for 30-60 minutes. Cells were re-pelleted by centrifugation, washed with PBS2%FBS and incubated with active rabbit anti-non-phosphorylated β-catenin antibody at room temperature for 60 minutes. Cells were re-pelleted by centrifugation, washed with PBS2%FBS and incubated with a conjugated secondary (donkey anti-rabbit – Alexa-647 – Thermo-Life tech, A31573) in the dark at room temperature for 30 minutes. Cells were re-pelleted by centrifugation, washed with PBS2%FBS and counterstained with 5µg of DAPI (1:1) before running through the flow cytometer. Unstained and single-stained (secondary antibody-only stained) cells were used as gaiting controls.

Unless specified otherwise, all data were collected on a BD FACS Canto II at the KU Leuven Flow Cytometry Core and analyzed using FlowJo v.10.6.2.

### Growth rate and Doubling time analysis

For Growth rate (µ) analysis, the following mathematical equation was used: 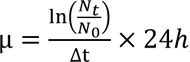, whereby N_0_ is the number of cells seeded, N_t_ is the number of cells harvested/recorded, and Δt is the hours of growth.

For Doubling time (t_d_) analysis, the following mathematical equation was used: 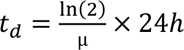.

### Conditioned media and co-culture analysis

Conditioned media (CM) was collected from TNBC cell lines recovering from chemotherapy treatment (5 days of treatment and 1 week of recovery) and filtered using a 0.45µM filter to ensure removing cell-debris. Filtered CM as concentrated 20x (20mL to 1mL) using Vivaspin centrifugal concentrator column with a molecular weight cutoff of 50kDa (Sigma Aldrich, Z614645-12EA). Filtered media was centrifuged for 45 minutes at 4°C. Concentrated CM was added to chemo-naïve TNBC cell lines for 48h in a 1:1 dilution (Concentrated CM:basal culture cancer media) and Wnt-activation levels were evaluated using FACS.

For co-culture experiments, MDA-MB-231 cell line was treated with either chemotherapeutic agent for 72h after which treatment was stopped and an equal number of chemo-naïve MDA-MB-231-TGP.mC cells was plated in the same dish and cultured in basal culture cancer media. After 72h of co-culture, Wnt-activation levels in the MDA-MB-231-TGP.mC cell line was evaluated using FACS.

### Cell line derived xenograft establishment and *in vivo* live imaging analysis

1 × 10^6^ MDA-MB-231 TOPFLASH cells were engrafted subcutaneously (1:1 PBS: growth-factor reduced Matrigel) into the right flank of female NMRI-Foxn1 mice (4-6 weeks old) to form a solid tumor. Upon observation of visible/palpable solid growth, tumor volumes were measured using digital calipers (and calculated as 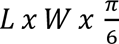; L: length and W: width). Animals were randomly assigned to one of three (or six) treatment groups (n = 7-8 mice per group) with an average tumor volume of 150mm^3^ per group. Docetaxel (15mg/kg) and Carboplatin (100mg/kg) were administered via intraperitoneal (IP) injection once weekly (1 cycle) for a total of three cycles. LGK-974 (2mg/kg) was administered daily via oral gavage for a total of 3 cycles (3 weeks). For assessment of Wnt-activation dynamics, animals were subjected to live-bioluminescent imaging before-, 24 h-, 48h-, and 72h-after chemotherapeutic administration. For live-bioluminescent imaging, animals were injected (IP) with the luciferase substrate D-luciferin (200µL of 15mg/ml - assuming an average animal weight of 24-26gr) (Perkin Elmer, 122799) and incubated for 10 minutes at room temperature before images were taken using IVIS Spectrum In Vivo Imaging System (Perkin Elmer). Wnt-activation signal was calculated as the bioluminescent signal captured by the IVIS Spectrum normalized to the tumor volume recorded per animal. Analysis of bioluminescent images was performed via the Aura software v.4.0.0. Tumor volume was recorded every 48h and body weight was closely monitored throughout the treatment course and recorded every 72-96h using an automatic scale. All animals were euthanized at the end of the treatment course and tumors (when available) were resected/collected for downstream analyses.

### RECIST Analysis

RECIST analysis was performed using tumor volumes measured and recorded (as described previously) at the onset of treatment and at the end of treatment (day of sacrifice). Relative tumor volume (RTV) was calculated by dividing the recorded volume at the end of treatment by the recorded volume at the onset of treatment. Response to therapy was based on the RECIST-based criterion: Complete response (CR), Partial response (PR), Stable disease (SD), and Progressive disease (PD); CR: RTV = 0, PR: 0 < RTV ≤ 0.657, SD: 0.657 < RTV ≤ 1.728, PD: RTV > 1.728.

### Tumorsphere Formation Assay

Cells were washed with PBS and collected/pelleted by centrifugation. Cells were resuspended in PBS4%FBS, counterstained with 5µg of DAPI (1:1) to eliminate dead cells before running through the SONY MA900 Multi-Application Cell Sorter. 1 × 10^3^single cells were sorted (based on their GFP expression) directly in ultra-low attachment 6-well plates (Fisher-Corning, 10154431) cultured in serum-free tumorsphere assay medium composed of DMEM/F12 (Gibco, 11320074), 1X B27 (Thermo-Scientific, 12587010), 10ng/mL basic fibroblast growth factor (bFGF) (Peprotech, 100-18b), 20ng/mL EGF (Peprotech, AF-100-15), and 2% growth-factor reduced Matrigel (Corning, 734-0268). Sorted cells were allowed fourteen days to grow, at the end of which, spheres were collected and centrifuged at 50g for 10minutes, resuspended gently, and transferred to 96-well plates (Fisher-Falcon,353072). Plates were briefly centrifuged at 50g for an additional 1 minutes to pull down larger spheroid (>60 µm) which were counted under a microscope (10X) using a tally counter.

### Next-Generation mRNA Sequencing

Total RNA was obtained from cells using the GenElute mammalian total RNA miniprep kit (Sigma, RTN350-1KT). RNA-sequencing (RNA-seq) libraries were prepared using 750 ng of total RNA using the KAPA stranded mRNAseq kit (Roche, 8098123702) according to the manufacturer’s specifications. 100 nM of KAPA-single index adapters (Roche, KK8702) were added to the A-tailed cDNA and the libraries underwent 10 cycles of amplification. Agentcourt AMPure XP beads (Beckman Coulter, A63880) were used for the 1X library clean-up. The fragment size of the libraries was assessed using the Agilent Bioanalyzer 2100 with the High Sensitivity DNA kit (Agilent, 5067-4626). The concentration of the libraries was measured by the High Sensitivity QuBit kit (Invitrogen, Q33230). Each library was diluted to 4 nM and pooled for single-end 50-bp sequencing on an Illumina Hiseq4000 with 20 – 27 million reads per sample (22 million reads on average).

### Bulk mRNA-sequencing analysis

FASTQ files generated from the sequencing (Sequencing run Fig. 1 and Fig. 3) were sent for downstream processing. Adapters were trimmed using Trimmomatic^83^ v0.39 and the trimmed FASTQ file was aligned to the GRCh38 genome (hg38) using the STAR aligner^84^ v2.7.10. Gene counts, gene annotation and sample read characteristics were obtained by applying standard filters within featureCounts^85^ from the subread package v2.0.3. Gene counts were then normalized using the variance stabilizing transformation (VST). Z-scores used to describe the gene expression distribution across samples were calculated using median absolute deviation whole the heatmaps comparing z-scores between samples were created using pheatmap v1.0.12. Differential gene expression analysis was performed using DESeq2^86^ and batch effects were accounted for in the (Wnt^High^ vs. Wnt^Low^ cohort). Volcano plots were created using EnhancedVolcano v1.18.0 using custom settings of FCcutoff = 0.6 and pCutoff = 0.05. Gene set variation analysis was performed using GSVA v1.48.3. Signature scores for the Caspase 3/Apoptosis^87^ and DTP**^Diap^**signatures (Supplementary Table 4) were calculated after the gene counts were transformed using both log2(x) +1 and VST methods. Box plots comparing the signature score(s) distribution between Wnt^High^ and Wnt^Low^ samples between treatment conditions were created using ggplot2 v3.4.3. Forest plots for regression analysis were created using forestplot v3.1.3. Analyses following the gene count extraction were all performed in R^88^ v4.3.0.

### Functional Enrichment Analysis of publicly available datasets

To identify sets of genes associated with a Wnt-active (Wnt^High^) signature, differential expression was performed on the TMM normalized gene counts using edgeR, taking into account treatment and Wnt population-status (High or Low) in the design matrix. The output this model was a list of differentially expressed genes. Wnt-signature, Diapause-DTP signature, and MYC-hallmark gene set enrichment was performed using the single-sample singscore R package.

### Patient derived organoid culture, treatment, and analysis

R1-IDC113 and R2-IDC159A PDO lines were gifted by our collaborator, Laboratory of Colinda Scheele – VIB-KU Leuven. Both PDO lines were maintained in growth factor reduced type 2 Cultrex (Biotechne/R&D Systems, 3533-010-02) with phenol red-free DMEM/F-12, HEPES (Gibco, 11039021) supplemented with 10mM Nicotinamide (Sigma/Merck, N0636-100G), 1.25mM N-acetyl-L-cystine (Sigma/Merck, A9165-5G), 500ng/mL Hydrocortisone, 100nM β-estradiol (E8875-250MG), 500nM SB202190 (Stem Cell Technologies, 72632), 500nM A83-01 (Stem Cell Technologies, 72022), 5uM Y-27632 (Stem Cell Technologies, 72304), 50µg/mL Primocin (Invivogen, ant-pm-05), 10µM Forskolin (Sigma/Merck, F3917-10MG), 1X B27 (50X – ThermoFisher Scientific, 17504044), 100ng/mL r-Noggin (Stem Cell Technologies, 78060), 5ng/mL FGF-10 (Stem Cell Technologies, 78037), 37.5 ng/mL Heregulin B-1 (Peprotech, 100-03), 5ng/mL EGF, 5ng/mL FGF-7 (Peprotech, 100-19), and 100µg/mL penicillin-streptomycin.

Both PDO lines were cultured in 24 well cell culture microplates – 22 mm x 20 (D x H) – at 37°C and 5% CO_2_ and maintained at 70-80% confluency. For passaging and plating, (ice-cold) 1X phosphate-buffered saline (Gibco, 10010-015) was used to wash and dissociate the BME followed by single cell enzymatic dissociation using 0.05% trypsin-EDTA (Gibco, 25200-056) and cell-pelleting by centrifugation for 5 minutes at 1500 rpm (4°C). Cells were counted manually via the BRAND counting chamber Neubauer improved (Sigma Aldrich/Merck, BR717810-1EA) under a 10X objective lens using a Leica DMi inverted microscope. The same microscope, equipped with a 2,5 Megapixel HD Microscope Camera Leica MC120 HC, was used to obtain images of cultured PDO-lines. Unless specified otherwise, both PDO models were plated according to the following seeding density: 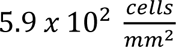

To determine working chemotherapy drug concentrations, PDO lines were treated with increasing concentrations of docetaxel (Taxotere, 0.0625–512 nM) and carboplatin (Carbosin, 0 – 1600 µM) for 96h. Cell metabolic activity, reflecting cell-number and –viability was assessed using the CellTiter-Glo® 3D Cell Viability Assay (Promega, G9682) and sigmoidal dose-response curves were generated to calculate the mean IC50 values of each drug that were used in the subsequent study. Chemotherapeutic agents were obtained from the pharmacy of Universitair Ziekenhuis (UZ) Leuven.

### Statistical Analysis

All data were analyzed using GraphPad Prism (v8.0.1), except for mRNA-sequencing derived data and transcriptomic datasets. Unless otherwise specified, comparisons between two groups were tested for statistical significance using Unpaired t-tests. Comparisons between multiple groups were performed using a One-way analysis of variance (ANOVA). Comparisons between multiple groups across multiple time points were performed using Two-way ANOVA. All statistical testing was corrected for multiple comparisons, using the Holm-Sidak method when comparing samples based on experimental design. For the reader’s convenience, all statistical tests and sample sizes are indicated in the figure legends.

For mRNA-sequencing derived data, regression analysis was performed to observe associations between outcomes (in-house gene signature scores/GSVA signature scores) and independent co-variate (Wnt^High^ vs. Wnt^Low^) per treatment condition (CAR or DOC or UNT) using lqmm v.1.5.8 and quantreg v5.97 while accounting for batch effects.

### Schematic Illustrations and artwork

All schematic illustrations were created using Biorender.com

## Data Availability

The bulk mRNA-sequencing data that support the findings of this study have been deposited in the Gene Expression Omnibus (GEO) repository under accession number GSE254558 (https://www.ncbi.nlm.nih.gov/geo/query/acc.cgi?acc=GSE254558).

For reviewers’ access, please enter the following token into the box to retrieve the dataset: **etsxyqgyrjkbhqt**

