## Supplementary Figures for "Chemotherapy-treated breast cancer cells activate the Wnt signaling pathway to enter a diapause-DTP state"

Supp. Fig. 1

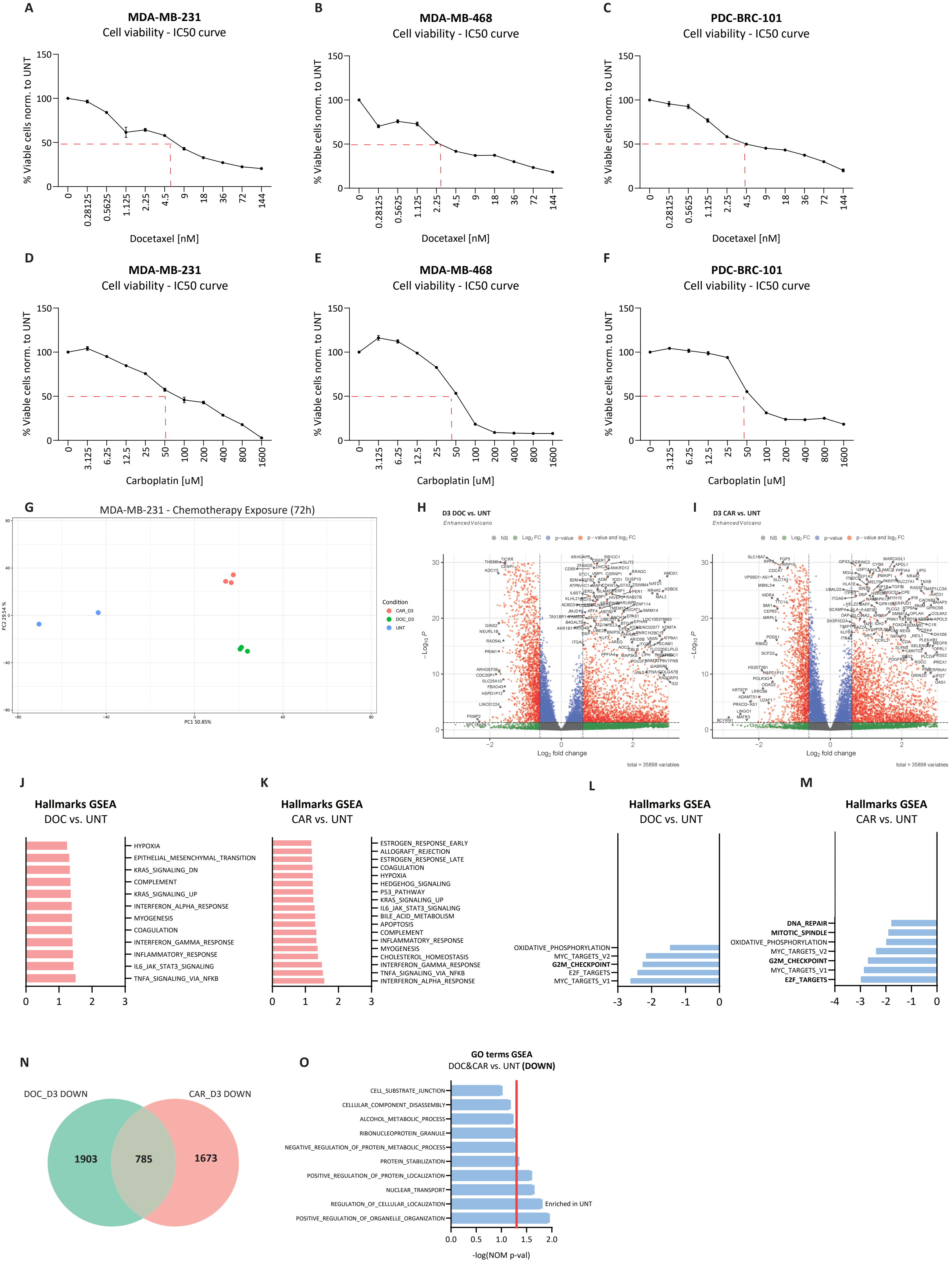
A
MDA-MB-231
Cell viability - IC50 curve
% Viable cells norm. to UNT
Docetaxel [nM]
0
0.28125
0.5625
1.125
2.25
4.5
9
18
36
72
144
B
MDA-MB-468
Cell viability - IC50 curve
C
PDC-BRC-101
Cell viability - IC50 curve
D
MDA-MB-231
Cell viability - IC50 curve
Carboplatin [uM]
0
3.125
6.25
12.5
25
50
100
200
400
800
1600
E
MDA-MB-468
Cell viability - IC50 curve
F
PDC-BRC-101
Cell viability - IC50 curve
G
MDA-MB-231 - Chemotherapy Exposure (72h)
PC1 50.85%
PC2 29.54 %
Condition
CAR_D3
DOC_D3
UNT
H
D3 DOC vs. UNT
EnhancedVolcano
I
D3 CAR vs. UNT
EnhancedVolcano
NS
Log2 FC
p−value
p−value and log2 FC
Log2 fold change
−Log10 P
total = 35898 variables
J
Hallmarks GSEA
DOC vs. UNT
HYPOXIA
EPITHELIAL_MESENCHYMAL_TRANSITION
KRAS_SIGNALING_DN
COMPLEMENT
KRAS_SIGNALING_UP
INTERFERON_ALPHA_RESPONSE
MYOGENESIS
COAGULATION
INTERFERON_GAMMA_RESPONSE
INFLAMMATORY_RESPONSE
IL6_JAK_STAT3_SIGNALING
TNFA_SIGNALING_VIA_NFKB
K
Hallmarks GSEA
CAR vs. UNT
ESTROGEN_RESPONSE_EARLY
ALLOGRAFT_REJECTION
ESTROGEN_RESPONSE_LATE
COAGULATION
HYPOXIA
HEDGEHOG_SIGNALING
P53_PATHWAY
KRAS_SIGNALING_UP
IL6_JAK_STAT3_SIGNALING
BILE_ACID_METABOLISM
APOPTOSIS
COMPLEMENT
INFLAMMATORY_RESPONSE
MYOGENESIS
CHOLESTEROL_HOMEOSTASIS
INTERFERON_GAMMA_RESPONSE
TNFA_SIGNALING_VIA_NFKB
INTERFERON_ALPHA_RESPONSE
L
Hallmarks GSEA
DOC vs. UNT
OXIDATIVE_PHOSPHORYLATION
MYC_TARGETS_V2
G2M_CHECKPOINT
E2F_TARGETS
MYC_TARGETS_V1
M
Hallmarks GSEA
CAR vs. UNT
DNA_REPAIR
MITOTIC_SPINDLE
OXIDATIVE_PHOSPHORYLATION
MYC_TARGETS_V2
G2M_CHECKPOINT
MYC_TARGETS_V1
E2F_TARGETS
N
DOC_D3 DOWN
CAR_D3 DOWN
1903
785
1673
O
GO terms GSEA
DOC&CAR vs. UNT (DOWN)
CELL_SUBSTRATE_JUNCTION
CELLULAR_COMPONENT_DISASSEMBLY
ALCOHOL_METABOLIC_PROCESS
RIBONUCLEOPROTEIN_GRANULE
NEGATIVE_REGULATION_OF_PROTEIN_METABOLIC_PROCESS
PROTEIN_STABILIZATION
POSITIVE_REGULATION_OF_PROTEIN_LOCALIZATION
NUCLEAR_TRANSPORT
REGULATION_OF_CELLULAR_LOCALIZATION
POSITIVE_REGULATION_OF_ORGANELLE_ORGANIZATION
Enriched in UNT
-log(NOM p-val)

**Supp. Fig. 1: Wnt transcriptional-activation precedes early drug-tolerant cell(s) enrichment upon chemotherapeutic treatment.**

**A-F)** Drug dose-response curves of MDA-MB-231, MDA-MB-468 and PDC-BRC-101 cell lines treated with increasing DOC (top) or CAR (bottom) for 72h. Dashed line represents 50% viability (normalized to UNT/basal culture conditions). **G)** Principal Component Analysis (PCA) plot of MDA-MB-231 cell line treated with DOC or CAR for 72h. **H-I)** Volcano plots displaying differentially regulated (down- (left) and up- (right) regulated) genes for MDA-MB-231 cell line treated with DOC (left) or CAR (right). Gene values are reported as $Log_2$FoldChange. **J-M)** De-regulated gene sets from Hallmarks database of DOC or CAR treatment (vs. UNT) analyzed by one-tailed GSEA ranked by Normalized Enrichment Score (NES), illustrating pathways and processes most significantly deregulated (up-regulated, pink & down-regulated, b) between chemo-treated and UNT-cells. **N)** Venn diagram indicating commonly downregulated (785) genes between DOC (1903) and CAR (1673) treatment (vs. UNT). **O)** Enriched gene sets from Gene Ontology (GO) terms databases analyzed by one-tailed GSEA ranked by a positive NES and based on commonly downregulated genes between DOC and CAR treatment. Red line indicates significance threshold value, (-log(NOM p-val) = 1.3). Data used to generate panels **G-O** was obtained from mRNA-sequencing of MDA-MB-231 cell line treated with DOC or CAR for 72h).

Supp. Fig. 2

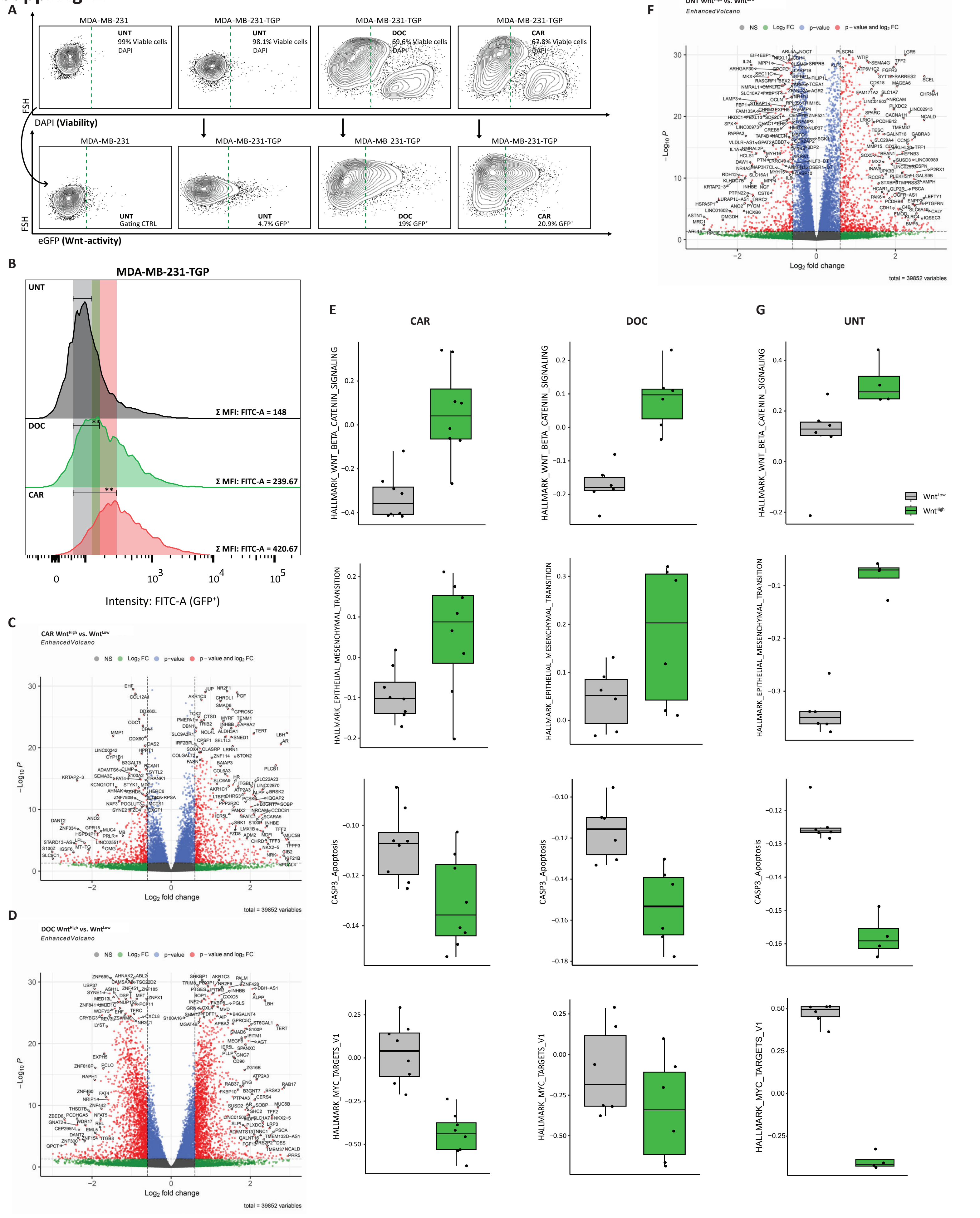
A
MDA-MB-231
MDA-MB-231-TGP
UNT
99% Viable cells
98.1% Viable cells
DOC
69.6% Viable cells
CAR
67.8% Viable cells
FSH
DAPI (Viability)
UNT
Gating CTRL
4.7% GFP+
19% GFP+
20.9% GFP+
eGFP (Wnt-activity)
B
MDA-MB-231-TGP
Σ MFI: FITC-A = 148
Σ MFI: FITC-A = 239.67
Σ MFI: FITC-A = 420.67
Intensity: FITC-A (GFP+)
C
CAR WntHigh vs. WntLow
EnhancedVolcano
NS
Log2 FC
p-value
p – value and log2 FC
Log2 fold change
total = 39852 variables
D
DOC WntHigh vs. WntLow
E
F
UNT WntHigh vs. WntLow
G
HALLMARK_WNT_BETA_CATENIN_SIGNALING
HALLMARK_EPITHELIAL_MESENCHYMAL_TRANSITION
CASP3_Apoptosis
HALLMARK_MYC_TARGETS_V1
WntLow
WntHigh

**Supp. Fig. 2: Parental and chemotherapy-treated Wnt$^{High}$ cells display DTP$^{Diap}$ cell-properties.**

**A)** Representative flow cytometry contour plots showing FACS gating strategy, displaying %Viable (DAPI$^{-}$) cells (left of the dashed green line – top) and % of Wnt$^{High}$ (GFP$^{+}$) (right of the dashed green line – bottom) MDA-MB-231-TGP cell line treated with DOC or CAR for 96h. **B)** An overlay of representative flow cytometry histograms representing the intensity of recorded GFP-expression (FITC-A channel) levels for MDA-MB-231-TGP TNBC cell line treated with DOC or CAR for 72h. Indicated numerical values denote the average median fluorescence intensity (MFI) recorded by the cytometer per sample. Multiple t-tests corrected for multiple comparisons using the Holms-Sidak method (n = 3 independent experiments) comparing MFI of DOC or CAR to UNT. **C-D)** Volcano plots displaying differentially regulated (down- (left) and up- (right) regulated) genes comparing sorted Wnt$^{High}$ vs. Wnt$^{Low}$ for MDA-MB-231-dTGFP cell line treated with CAR or DOC. Gene values are reported as $Log_2$FoldChange. Dot colors are clearly indicated in the plot. **E)** Box plots depicting the marked differences in absolute scores of selected gene signatures between sorted Wnt$^{High}$ and Wnt$^{Low}$ cells across CAR- and DOC-treated samples. Batch effects were not accounted for in this analysis. **F)** Volcano plot displaying differentially regulated (down- (left) and up- (right) regulated) genes comparing sorted Wnt$^{High}$ vs. Wnt$^{Low}$ for MDA-MB-231-dTGFP cell line in UNT conditions. Gene values are reported as $Log_2$FoldChange. Dot colors are clearly indicated in the plot. **G)** Box plots depicting the marked differences in absolute scores of selected gene signatures between sorted Wnt$^{High}$ and Wnt$^{Low}$ cells in UNT conditions. Batch effects were not accounted for in this analysis. p values are indicated as *$p < 0.05$, **$p < 0.01$, ***$p < 0.001$, ****$p < 0.0001$, and ns, not significant.

**Supp. Fig. 3**

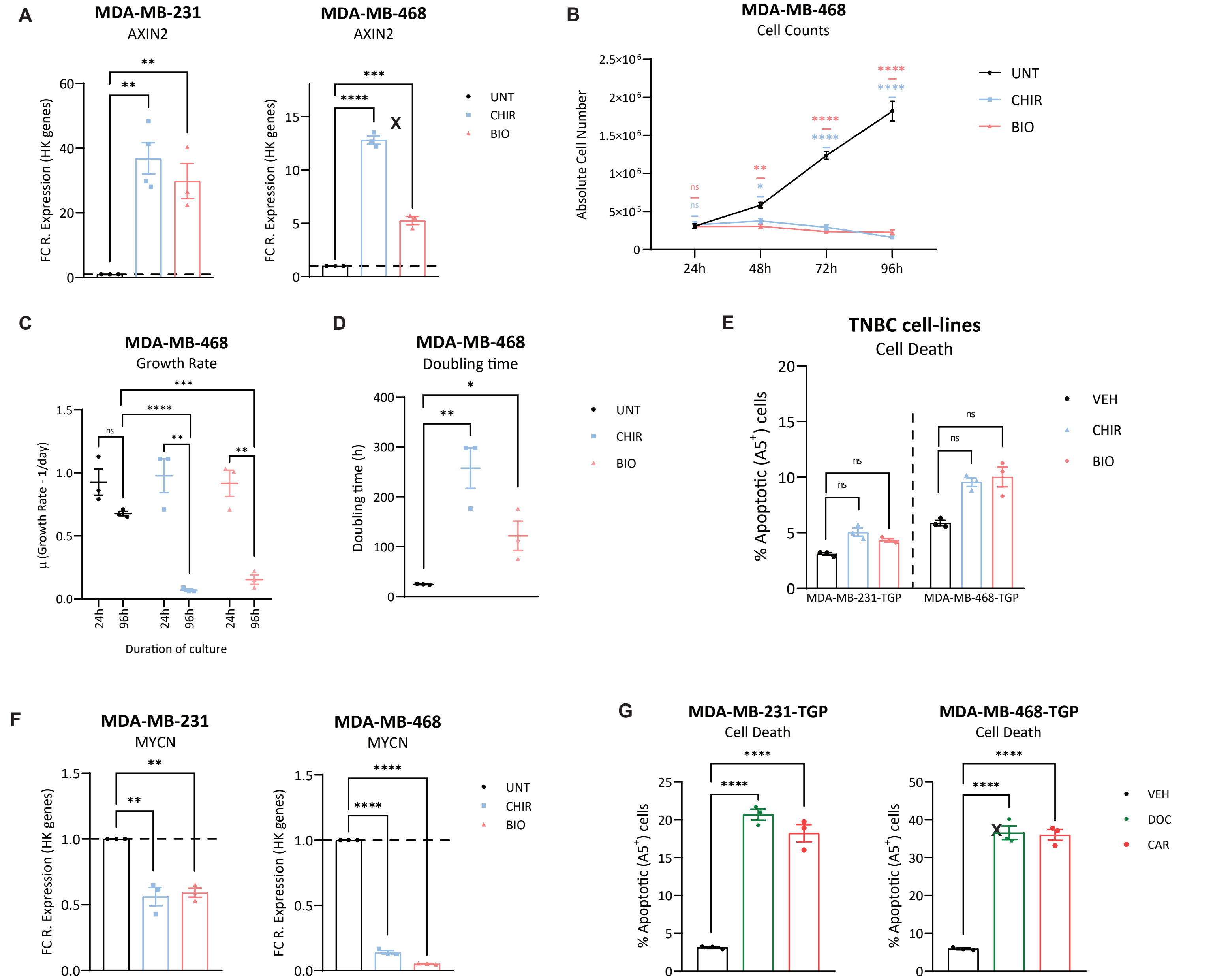


**Supp. Fig. 3: Wnt pathway-activation is sufficient to induce a DTP$^{Diap}$ state in parental TNBC cells.**

**A)** Gene expression levels obtained via RT-qPCR of Wnt-target (AXIN2) for MDA-MB-231 and MDA-MB-468 cell lines treated with CHIR or BIO for 72h, displayed as fold change (to UNT) of $2^{-dCt}$ values (relative to HK-genes). Unpaired t-tests based on relative expression values ($2^{-dCt}$) (n = 3 independent experiments). Data are presented as Mean ± SEM. **B)** Absolute cell number of MDA-MB-231 cell line treated with CHIR or BIO for a time-course of 96h. Two-way ANOVA corrected for multiple comparisons using Tukey's test (n = 3 independent experiments). Data are presented as Mean ± SEM. **C)** Growth rate of MDA-MB-468 cell line at 24h and 96h under UNT or under CHIR and BIO treatment conditions. Multiple t-tests corrected for multiple comparisons using the Holms-Sidak method (n = 3 independent experiments). Data are represented as Mean ± SEM. **D)** Doubling time of MDA-MB-468 cell line at 96h under UNT or under CHIR and BIO treatment conditions. Multiple t-tests corrected for multiple comparisons using the Holms-Sidak method (n = 3 independent experiments). Data are represented as Mean ± SEM. **E)** Flow cytometry analysis displaying % of Annexin $V^+$ cells of MDA-MB-231-TGP and MDA-MB-468-TGP cell lines treated with CHIR or BIO for 72h (sole treatments). One-way ANOVA corrected for multiple comparisons using the Dunnett method (n = 3 independent experiments). Data are presented as Mean ± SEM. **F)** Gene expression levels obtained via RT-qPCR of MYCN for MDA-MB-231 and MDA-MB-468 cell lines treated with CHIR or BIO for 72h, displayed as fold change (to UNT) of $2^{-dCt}$ values (relative to HK-genes). Unpaired t-tests based on relative expression values ($2^{-dCt}$) (n = 3 independent experiments). Data are presented as Mean ± SEM. **G)** Flow cytometry analysis displaying % of Annexin $V^+$ cells of MDA-MB-231-TGP (left) and MDA-MB-468-TGP (right) cell lines treated with chemotherapy (DOC or CAR) for 72h (sole treatments). One-way ANOVA corrected for multiple comparisons using the Dunnett method (n = 3 independent experiments). Data are presented as Mean ± SEM. p values are indicated as *p < 0.05, **p < 0.01, ***p < 0.001, ****p < 0.0001, and ns, not significant.

**Supp. Fig. 4**

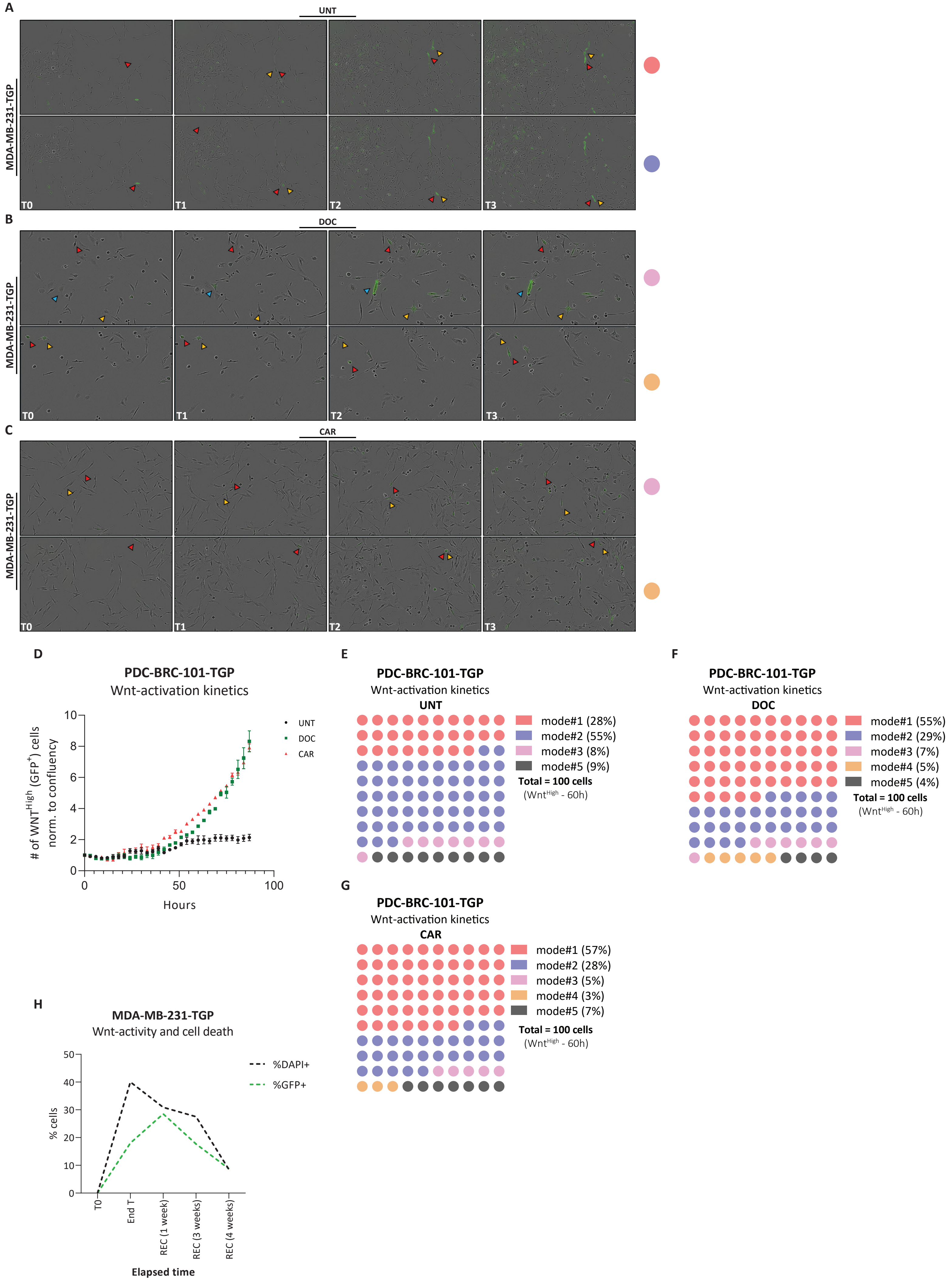
A
UNT
MDA-MB-231-TGP
T0
T1
T2
T3
B
DOC
MDA-MB-231-TGP
T0
T1
T2
T3
C
CAR
MDA-MB-231-TGP
T0
T1
T2
T3
D
PDC-BRC-101-TGP
Wnt-activation kinetics
### of WNT^High (GFP^+) cells norm. to confluency
Hours
0
50
100
0
2
4
6
8
10
UNT
DOC
CAR
E
PDC-BRC-101-TGP
Wnt-activation kinetics
UNT
mode#1 (28%)
mode#2 (55%)
mode#3 (8%)
mode#5 (9%)
Total = 100 cells
(Wnt^High - 60h)
F
PDC-BRC-101-TGP
Wnt-activation kinetics
DOC
mode#1 (55%)
mode#2 (29%)
mode#3 (7%)
mode#4 (5%)
mode#5 (4%)
Total = 100 cells
(Wnt^High - 60h)
G
PDC-BRC-101-TGP
Wnt-activation kinetics
CAR
mode#1 (57%)
mode#2 (28%)
mode#3 (5%)
mode#4 (3%)
mode#5 (7%)
Total = 100 cells
(Wnt^High - 60h)
H
MDA-MB-231-TGP
Wnt-activity and cell death
% cells
0
10
20
30
40
50
%DAPI+
%GFP+
T0
End T
REC (1 week)
REC (3 weeks)
REC (4 weeks)
Elapsed time

**Supp. Fig. 4: Induction of transient *de novo* Wnt signaling transcriptional-activation in response to chemotherapy in TNBC cell lines.**

**A-C)** Snapshots of still-frames from time-lapse live imaging experiments of MDA-MB-231-TGP TNBC cell line UNT (bottom), treated with DOC (middle), or CAR (bottom) presenting mode #1, #2, #3, and #4 (color coding in scheme shown in **Fig. 4B**. $T_0$ indicates 0h, $T_1$ indicates 30h, $T_2$ indicates 48h, and $T_3$ indicates 60h. Yellow, red, and blue arrows indicate the same cell followed over the treatment period spanning different images (horizontal). **D)** Number of $Wnt^{High}$ ($GFP^+$) cells detected by live-cell imaging normalized to the confluency of the well (total number of cells recorded) for PDC-BRC-101-TGP cell line treated with DOC or CAR. **E-G)** Live-imaging quantification of different possible mechanisms of chemotherapy-induced Wnt-activation in PDC-BRC-101-TGP TNBC cell line UNT or treated with DOC or CAR. n = 100 cells tracked every 2h for 60h, per treatment condition. Every single cell tracked is represented as a circle and color coded with the scheme shown in **Fig. 4B**. **H)** Flow Cytometry analysis displaying % of $Wnt^{High}$ ($GFP^+$) and % of $DAPI^+$ cells of MDA-MB-231-TGP cell line treated with DOC following the treatment scheme shown in **Fig. 4H**. p values are indicated as *$p < 0.05$, **$p < 0.01$, ***$p < 0.001$, ****$p < 0.0001$, and ns, not significant.

### Supp. Fig. 5

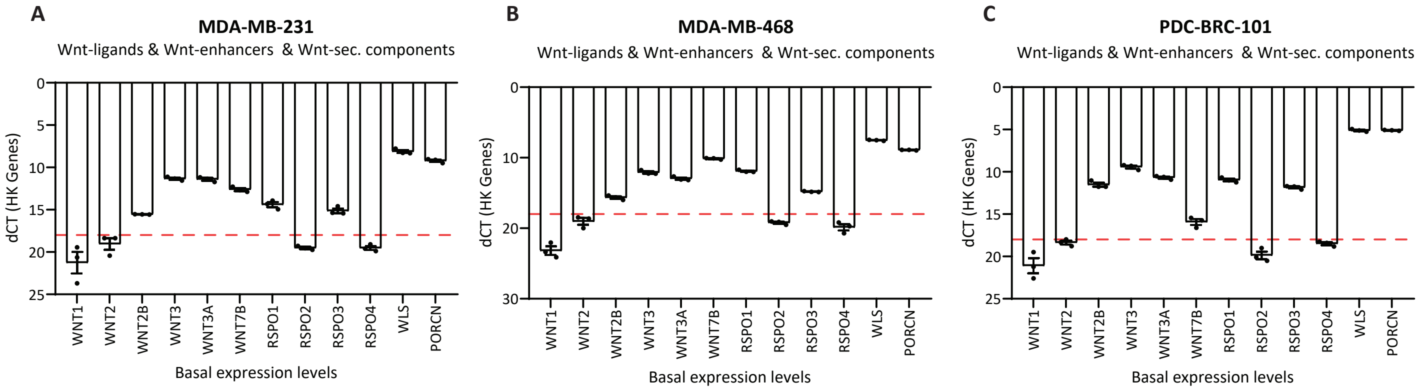


**Supp. Fig. 5: Chemotherapeutic treatment induces elevated transcriptional expression of Wnt ligands, Wnt enhancers, and Wnt secretion machinery components.**
**A-C)** Gene expression levels obtained via RT-qPCR of various Wnt ligands, Wnt enhancers, and Wnt secretion machinery components for MDA-MB-231, MDA-MB-468, and PDC-BRC-101 cell lines in basal culture conditions displayed as dCt values (relative to HK-genes) (n = 3 independent experiments). The dashed red line represents a threshold dCt value of 18. Genes that record dCt values above 18 are not considered robustly expressed and are therefore not subject to further analysis in our study.

Supp. Fig. 6

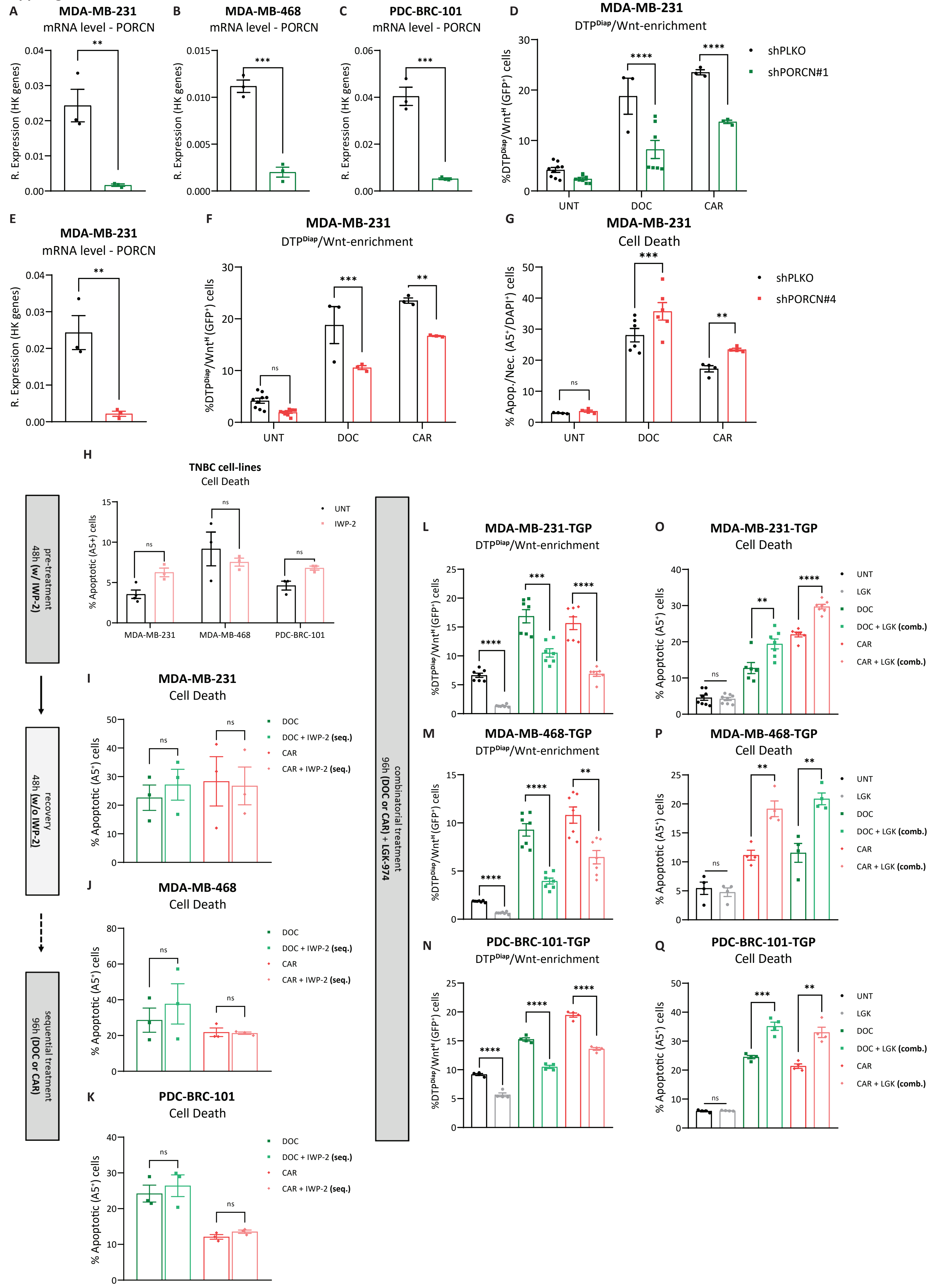
A
MDA-MB-231
mRNA level - PORCN
R. Expression (HK genes)
B
MDA-MB-468
mRNA level - PORCN
C
PDC-BRC-101
mRNA level - PORCN
D
MDA-MB-231
DTP^Diap/Wnt-enrichment
%DTP^Diap/Wnt^H (GFP^+) cells
shPLKO
shPORCN#1
UNT
DOC
CAR
E
MDA-MB-231
mRNA level - PORCN
F
MDA-MB-231
DTP^Diap/Wnt-enrichment
G
MDA-MB-231
Cell Death
% Apop./Nec. (A5^+/DAPI^+) cells
shPLKO
shPORCN#4
H
TNBC cell-lines
Cell Death
% Apoptotic (A5+) cells
UNT
IWP-2
MDA-MB-231
MDA-MB-468
PDC-BRC-101
pre-treatment
48h (w/ IWP-2)
recovery
48h (w/o IWP-2)
sequential treatment
96h (DOC or CAR)
I
MDA-MB-231
Cell Death
% Apoptotic (A5^+) cells
DOC
DOC + IWP-2 (seq.)
CAR
CAR + IWP-2 (seq.)
J
MDA-MB-468
Cell Death
K
PDC-BRC-101
Cell Death
combinatorial treatment
96h (DOC or CAR) + LGK-974
L
MDA-MB-231-TGP
DTP^Diap/Wnt-enrichment
M
MDA-MB-468-TGP
DTP^Diap/Wnt-enrichment
N
PDC-BRC-101-TGP
DTP^Diap/Wnt-enrichment
O
MDA-MB-231-TGP
Cell Death
UNT
LGK
DOC
DOC + LGK (comb.)
CAR
CAR + LGK (comb.)
P
MDA-MB-468-TGP
Cell Death
Q
PDC-BRC-101-TGP
Cell Death
ns

**Supp. Fig. 6: Wnt ligand secretion-inhibition hinders $DTP^{Diap}/Wnt^{H}$ population enrichment.**

**A-C)** Gene expression levels obtained via RT-qPCR of PORCN of MDA-MB-231, MDA-MB-468, and PDC-BRC-101 (shPLKO vs. shPORCN#1) cell lines in basal conditions. Unpaired t-tests based on relative expression values ($2^{-dCt}$) (n = 3 independent experiments). Data are presented as Mean ± SEM. **D)** Flow cytometry analysis displaying % of $DTP^{Diap}/Wnt^{H}$ ($GFP^{+}$) cells of MDA-MB-231 (shPLKO vs. shPORCN#1) cell line treated with DOC or CAR for 96h. Multiple t-tests corrected for multiple comparisons using the Holms-Sidak method (n = 3,4 independent experiments). Data are presented as Mean ± SEM. **E)** Gene expression levels obtained via RT-qPCR of PORCN of MDA-MB-231 (shPLKO vs. shPORCN#4) cell line in basal culture conditions. Unpaired t-tests based on relative expression values ($2^{-dCt}$) (n = 3 independent experiments). Data are presented as Mean ± SEM. **F)** Flow cytometry analysis displaying % of $DTP^{Diap}/Wnt^{H}$ ($GFP^{+}$) cells of MDA-MB-231 (shPLKO vs. shPORCN#4) cell line treated with DOC or CAR for 96h. Multiple t-tests corrected for multiple comparisons using the Holms-Sidak method (n = 3,4 independent experiments). Data are presented as Mean ± SEM. **G)** Flow cytometry analysis displaying % of Apoptotic and Necrotic (Annexin $V^{+}$ and $DAPI^{+}$) cells of MDA-MB-231 (shPLKO vs. shPORCN#4) cell line treated with DOC or CAR for 96h. Multiple t-tests corrected for multiple comparisons using the Holms-Sidak method (n = 4 independent experiments). Data are presented as Mean ± SEM. **H)** Flow cytometry analysis displaying % of Apoptotic (Annexin $V^{+}$) cells of TNBC cell lines pre-treated with IWP-2 (10μM) for 48h. Multiple t-tests corrected for multiple comparisons using the Holms-Sidak method (n = 3 independent experiments). Data are presented as Mean ± SEM. **I-K)** Flow cytometry analysis displaying % of Apoptotic (Annexin $V^{+}$) cells of TNBC-TGP cell lines treated with DOC or CAR for 96h (with or without IWP-2 pre-treatment). Multiple t-tests corrected for multiple comparisons using the Holms-Sidak method (n = 3 independent experiments). Data are presented as Mean ± SEM. **L-N)** Flow cytometry analysis displaying % of $DTP^{Diap}/Wnt^{H}$ ($GFP^{+}$) cells of MDA-MB-231-TGP, MDA-MB-468-TGP, and PDC-BRC-101-TGP cell lines treated with DOC or CAR for 96h (sole or in combination with LGK-974, 2μM). Multiple t-tests corrected for multiple comparisons using the Holms-Sidak method (n = 4 independent experiments). Data are presented as Mean ± SEM. **O-Q)** Flow cytometry analysis displaying % of Apoptotic (Annexin $V^{+}$) cells of TNBC-TGP cell lines treated with DOC or CAR for 96h (sole or in combination with LGK-974). Multiple t-tests corrected for multiple comparisons using the Holms-Sidak method (n = 4 independent experiments). Data are presented as Mean ± SEM. p values are indicated as $^{*}p < 0.05$, $^{**}p < 0.01$, $^{***}p < 0.001$, $^{****}p < 0.0001$, and ns, not significant.

Supp. Fig. 7

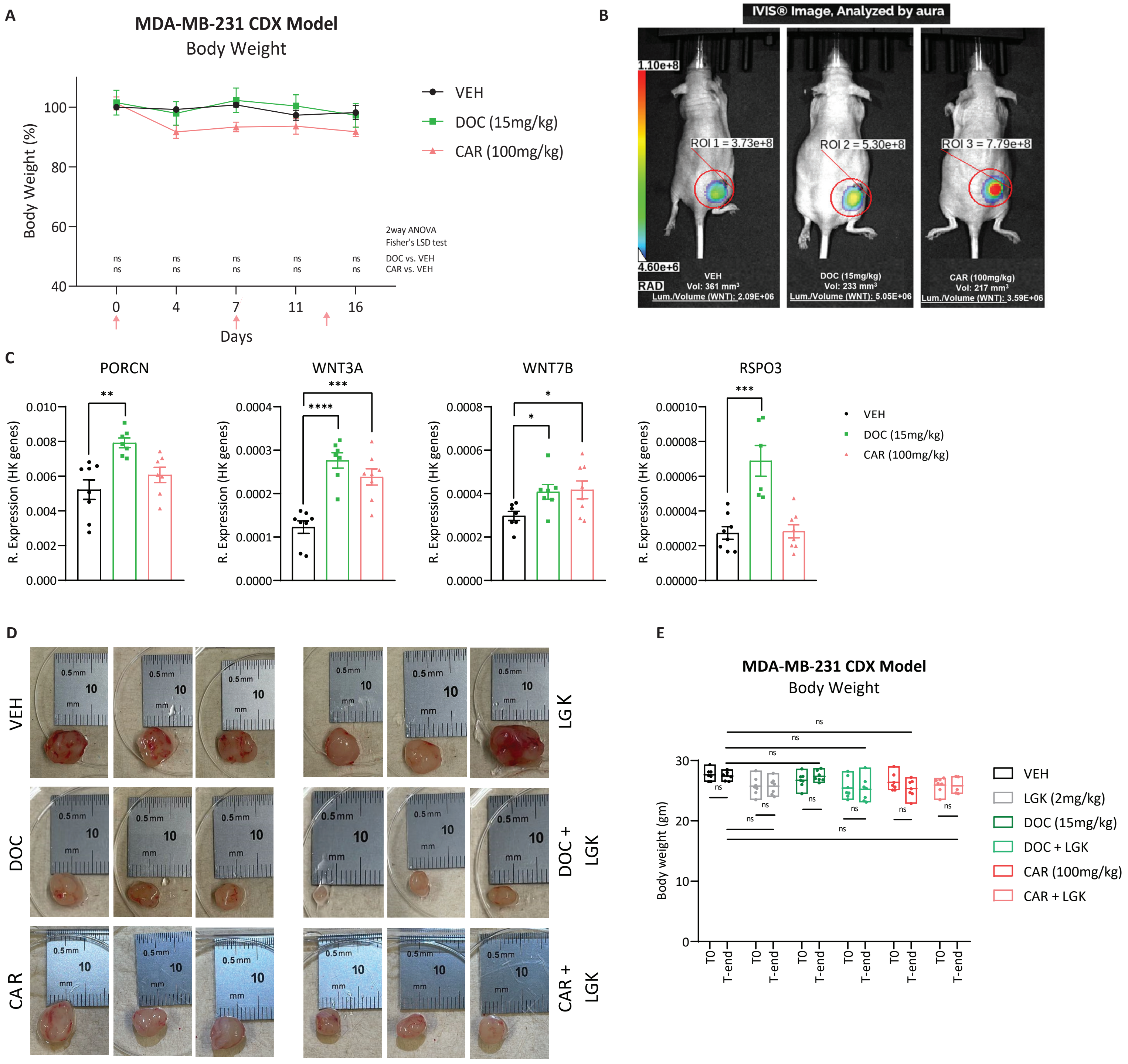


**Supp. Fig. 7 : Inhibition of Wnt ligand-secretion and chemotherapeutic treatment synergistically sensitize *in vivo* xenograft TNBC model to treatment.**
**A)** % of bodyweight change of initial treatment bodyweight of xenograft models treated with VEH, DOC, or CAR. Two-way ANOVA with Fisher's LSD test (n = 7 mice for all treatment groups). Data are presented as Mean ± SEM. **B)** Representative bioluminescent image obtained via IVIS Spectrum displaying one mouse from each of the three treatment groups (VEH – left, DOC – middle, and CAR – right) with supplementary information regarding the tumor volume of each mouse and the calculated Wnt-activation signal. **C)** Gene expression levels obtained via RT-qPCR of Wnt-activators in samples resected from xenograft models treated with VEH, DOC, or CAR. Unpaired t-tests based on relative expression values ($2^{-dCt}$) (n = 7 mice for all treatment groups). Data are presented as Mean ± SEM. **D)** Representative images (3 per treatment group) of resected tumors from xenograft models treated with VEH, LGK, DOC, DOC+LGK, CAR, and CAR+LGK. **E)** Bodyweight measurements evaluated at T0 (onset of treatment) and T-end (day of sacrifice) of xenograft models treated with VEH, LGK, DOC, DOC+LGK, CAR, and CAR+LGK. Two-way ANOVA test corrected for multiple comparisons using Tukey's method (n = 8,7,5 mice per treatment group). Data are presented as Mean ± SEM. p values are indicated as *p < 0.05, **p < 0.01, ***p < 0.001, ****p < 0.0001, and ns, not significant.

#### Supp. Fig. 8

A

| Sample ID | Tumor | Treatment Status | Treatment | Response | EUSOMA |
|---|---|---|---|---|---|
| R1-IDC113 | Primary Breast | Pretreated | ddAC | Progression | EUSOMA 3 |
| R2-IDC159A | Primary Breast | Pretreated | AC + paclitaxel + carboplatin + fulvestrant + radiotherapy | Progression | EUSOMA 3 |

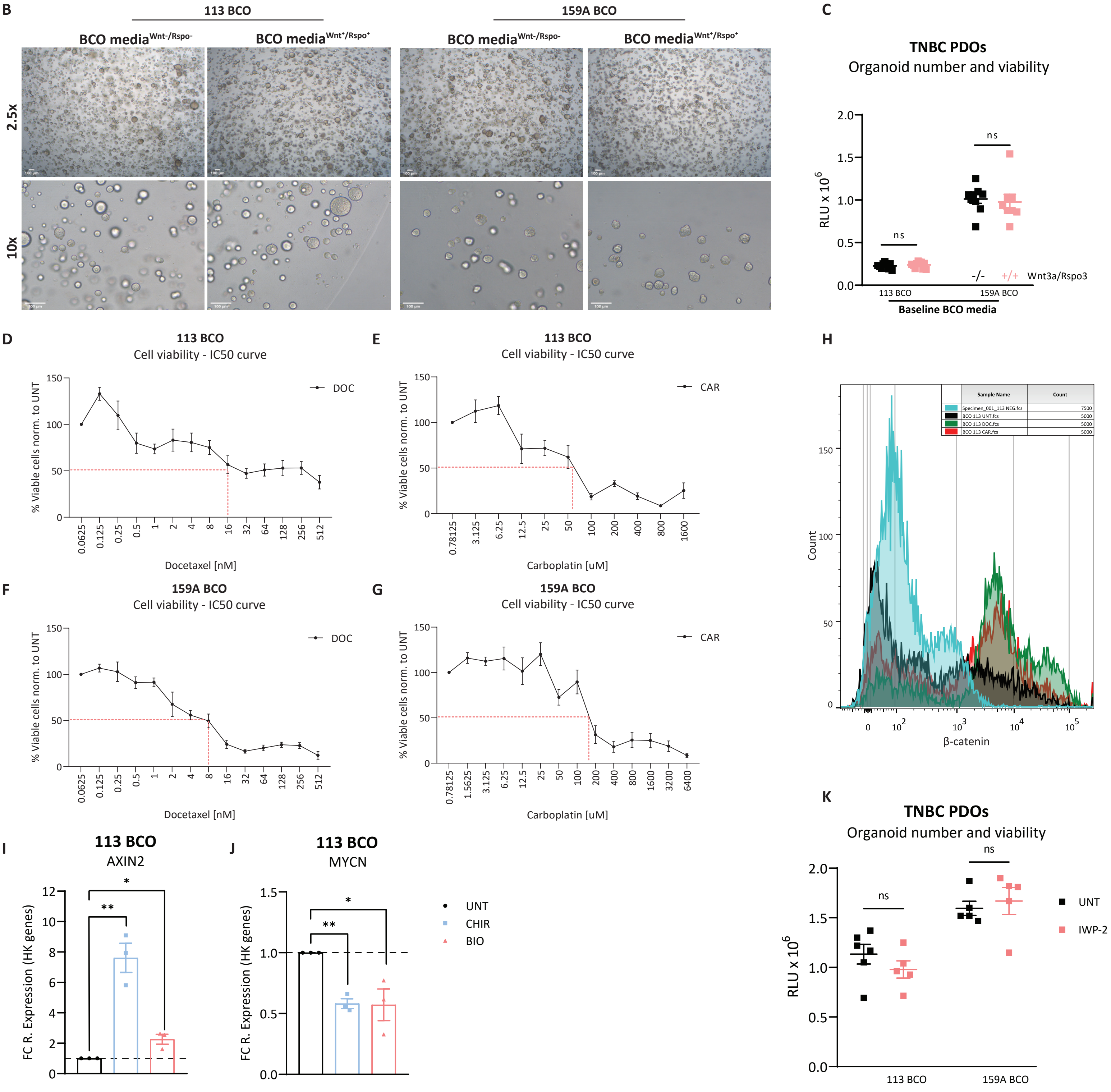


**Supp. Fig. 8: Preclinical PDO models recapitulate chemotherapy-mediated Wnt-activation and sensitization to synergistic Wnt ligand secretion-inhibition.** **A)** Overview of supplementary information of the TNBC PDO models used in the study. **B)** Phase-contrast images of TNBC-PDO models, 113 BCO (left) and 159A BCO (right) cultured in media not-supplemented with Wnt-Rspo (BCO media$^{Wnt-/Rspo-}$) and media supplemented with Wnt-Rspo (BCO media$^{Wnt+/Rspo+}$) at 2.5x (top) and 10x (bottom) magnification. **C)** Number and viability levels of 113 BCO (left) and 159A BCO (right) models cultured in media not-supplemented with Wnt-Rspo (BCO media$^{-/-}$) and media supplemented with Wnt-Rspo (BCO media$^{+/+}$). Unpaired t-tests (n = 3 independent experiments). Data are presented as Mean ± SEM. **D-G)** Drug dose-response curves of 113 and 159A BCO model treated with increasing DOC or CAR for 96h. Dashed line represents 50% viability (normalized to UNT/basal culture conditions). **H)** Representative histograms obtained from flow cytometry-based β-catenin staining displaying counts and intensity of β-catenin signal from 113 BCO model treated with DOC or CAR for 96h. Light blue signal displays background signal obtained from an unstained (NEG) sample. **I-J)** Gene expression levels obtained via RT-qPCR of Wnt-target gene (AXIN2) and MYCN for 113 BCO model treated with CHIR or BIO for 72h, displayed as fold change (to UNT) of $2^{-dCt}$ values (relative to HK-genes). Unpaired t-tests (n=3 independent experiments). Data are represented as Mean ± SEM. **K)** Number and viability levels of 113 BCO (left) and 159A BCO (right) models cultured in basal conditions and treated with IWP-2 (50µM) for 96h. Unpaired t-tests (n = 3 independent experiments). Data are presented as Mean ± SEM. p values are indicated as *p < 0.05, **p < 0.01, ***p < 0.001, ****p < 0.0001, and ns, not significant.

**Supplementary Videos**
**SV1 and SV2**: Time-lapse video of MDA-MB-231-TGP cell line in UNT (basal culture conditions) for 48 hours. Images taken every 2 hours under 10x objective lens.
**SV3 and SV4**: Time-lapse video of MDA-MB-231-TGP cell line treated with DOC for 48 hours. Images taken every 2 hours under 10x objective lens.
**SV5 and SV6**: Time-lapse video of MDA-MB-231-TGP cell line treated with CAR for 48 hours. Images taken every 2 hours under 10x objective lens.
